# Structural and Computational Design of a SARS-2 Spike Antigen with Increased Receptor Binding Domain Exposure and Improved Immunogenicity

**DOI:** 10.1101/2022.11.29.518231

**Authors:** James A. Williams, Marco Biancucci, Laura Lessen, Sai Tian, Ankita Balsaraf, Lynn Chen, Chelsy Chesterman, Giulietta Maruggi, Sarah Vandepaer, Ying Huang, Corey P. Mallett, Ann-Muriel Steff, Matthew James Bottomley, Enrico Malito, Newton Wahome, Wayne D. Harshbarger

## Abstract

Emerging SARS-CoV-2 variants of concern challenge the efficacy of approved vaccines and emphasize the need for improved antigens. Using an evolutionary-based design approach starting from the widely used engineered Spike antigen, S-2P, we sought to increase antigen production levels and the exposure of highly conserved and neutralization sensitive receptor-binding domain (RBD) epitopes. Thirty-six prototypes were generated *in silico*, of which fifteen were produced and tested in biochemical assays. Design S2D14, which contains 20 mutations within the Spike S2 domain, showed a 6-fold increase in expression while preserving similar thermal stability and antigenicity as S-2P. Cryo-EM structures indicate that the dominant populations of S2D14 particles have RBDs in exposed states, and analysis of these structures revealed how modifications within the S2 domain balance trimer stability and RBD accessibility through formation and removal of hydrogen bonds and surface charge alterations. Importantly, vaccination of mice with adjuvanted S2D14 resulted in higher levels of neutralizing antibodies than adjuvanted S-2P against SARS-CoV-2 Wuhan strain and four variants of concern. These results can guide the design of next generation vaccines to combat current, and future coronaviruses and the approaches used may be broadly applicable to streamline the successful design of vaccine antigens.

## Introduction

Severe acute respiratory syndrome coronavirus 2 (SARS-CoV-2), the causative agent for coronavirus disease 2019 (COVID-19), was first identified in Wuhan, China before spreading globally and being declared a pandemic in March 2020. With over 613 million confirmed cases globally, resulting in over 6.5 million deaths, COVID-19 remains a significant global health burden. Vaccines, such as mRNA-1273 (Moderna), BNT162b2 (Pfizer/Biontech), and Ad.26.COV2.S (Janssen), are all based on the sequence of the original Wuhan-Hu-1 strain of the Spike fusion glycoprotein (S), engineered to remain in the prefusion state, which is the primary target of neutralizing antibodies (nAbs). However, the efficacies of these first-generation vaccines are diminished against newly circulating variants of concern (VoCs) such as Alpha (B.1.1.7), Beta (B.1.351), Delta (B.1.617.2), and Omicron (BA.1, BA.2, BA.4/5, XBB, and BQ.1.1), which evade neutralizing antibodies and/or cell mediated immunity due to mutations in the S protein [1-8]. Though booster vaccines have been developed to match S protein sequences of circulating variants [9-12], there is no guarantee that these updated vaccines will protect against future strains of the virus [7, 8, 13-15]. Therefore, there is a need for the development of vaccine antigens which can elicit Abs that are more efficient in neutralizing future variants, thus providing broader and/or, longer lasting protection.

The S glycoprotein is composed of S1 and S2 subunits that mediate host cell attachment to initiate virus-cell fusion [16, 17]. The S1 subunit comprises the N-terminal domain (NTD) and the receptor-binding domain (RBD) with subdomains SD1 and SD2 undergoing hinge-like motions that promote RBD movement between closed and opened conformations [18, 19]. Engagement of the open RBD conformation with human angiotensin converting enzyme 2 receptor (hACE2) triggers a large-scale conformational rearrangement in the S2 domain, which contains the fusion machinery necessary to mediate viral fusion and infectivity [17, 20]. A large portion of the nAb response elicited during natural infection or vaccination is directed towards epitopes spanning the RBD [21-24]. Epitopes in less accessible regions of the RBD are more conserved among circulating coronaviruses, consistent with Abs targeting these regions being able to cross-neutralize other coronaviruses [21, 25, 26]. Several groups have developed adjuvanted RBD only vaccines based on monomeric or multivalent display of the RBD, with several human phase I/II and phase III trials showing a safe and immunogenic response, thus making the RBD, or S in the RBD-open state, an attractive vaccine antigen [27-37].

Engineering of the S protein to remain in the prefusion state was accomplished through rational, structure-based design and the insertion of two proline residues, K986P and V987P (named S-2P), which are located between the heptad repeat region 1 (HR1) and the central helix domain (CH) [20]. S-2P was later modified to have higher thermostability and greater protein expression through incorporation of additional prolines (F817P, A892P, A899P, A942P) (referred to as S-6P or HexaPro), which are located between the fusion peptide proximal region, HR1, and the CH domain [38]. Similar structure-based approaches have also been instrumental in the design of vaccine antigens for other pathogens such as the respiratory syncytial virus fusion protein F, metapneumovirus fusion protein F, influenza HA, and HIV-1 Envelope [39-44]. However, the identification of an antigen with the preferred qualities (i.e., conformation, stability, expression) often requires testing of hundreds of single-point mutations followed by combinatorial design which can lead to many failures.

Data-driven computational approaches have the potential to reveal mutable space within a protein’s structure that may not be obvious in a rational-based setting and assist antigen design by identifying sequences that yield desired protein characteristics [45-49]. One such computational method is PROSS (Protein Repair One-Stop Shop), which is a design algorithm using evolutionary information in multi-sequence alignments to focus the protein design search on residues that are functional in nature [45]. Combined with the Rosetta design suite and energy scoring function [50, 51], false positive predictions can be minimized and the number of variants that must be experimentally tested are reduced [45, 52, 53]. Success has been shown for multiple enzymes and the HIV gp140 glycoprotein, where utilization of PROSS led to enhanced protein stability and greater expression of functional protein compared to wild-type [45, 53, 54].

Here, we adapted the PROSS workflow to perform design based on multi-sequence alignment of related coronavirus glycoproteins, and incorporated symmetry protocols to uniformly mutate the S protein’s trimeric structure. The evolutionary consensus design strategy resulted in a novel prefusion S antigen that had biochemical and biophysical characteristics comparable to S-2P, but with greater expression and capable of eliciting higher levels of nAbs against Wuhan and VoCs in mice. High resolution structures by cryo-EM reveal that a potential mechanism of action for the improved quality of the Ab response was achieved through stabilization of the RBD-open conformation. This work increases our understanding of the molecular determinants leading to improved S antigens and will be valuable for informing the design of spike-based vaccines which can elicit broad protection against emerging and future strains of the virus.

## Results

### Computational design of SARS-CoV-2 Spike antigens

The evolutionary consensus design workflow combines three distinct steps, as previously described [45]. First, we generated a multi-sequence alignment of 500 non-redundant Spike protein sequences from various betacoronavirus lineages (lineages *A-D)* which we obtained from the BLAST database [55] (**Data S1**), allowing for the identification of residues with natural variation. Next, Rosetta atomistic design simulations [51] were used to curate which set(s) of single point mutations could be applied to the above identified residues to obtain novel spike designs with predicted lower free energy (i.e., improved stability) compared to the target S-2P antigen sequence [20]. To generate an initial S antigen model, we chose to incorporate a symmetry-based protocol [56] with a molecular structure that has all three RBDs in the open conformation (**Fig. 1**). Additionally, our starting model contained the S-2P di-proline mutations to ensure that the prefusion conformation of Spike was maintained [20], as well as the mutation D614G, located within the SD2 domain, which was an over-represented mutation in the above sequence alignments for circulating strains at the time of this study [57, 58]. In the final step, Rosetta combinatorial sequence optimization was used to generate constructs with energy profiles more favorable than the initial S antigen model (**Fig. S1**).

**Fig. 1.**
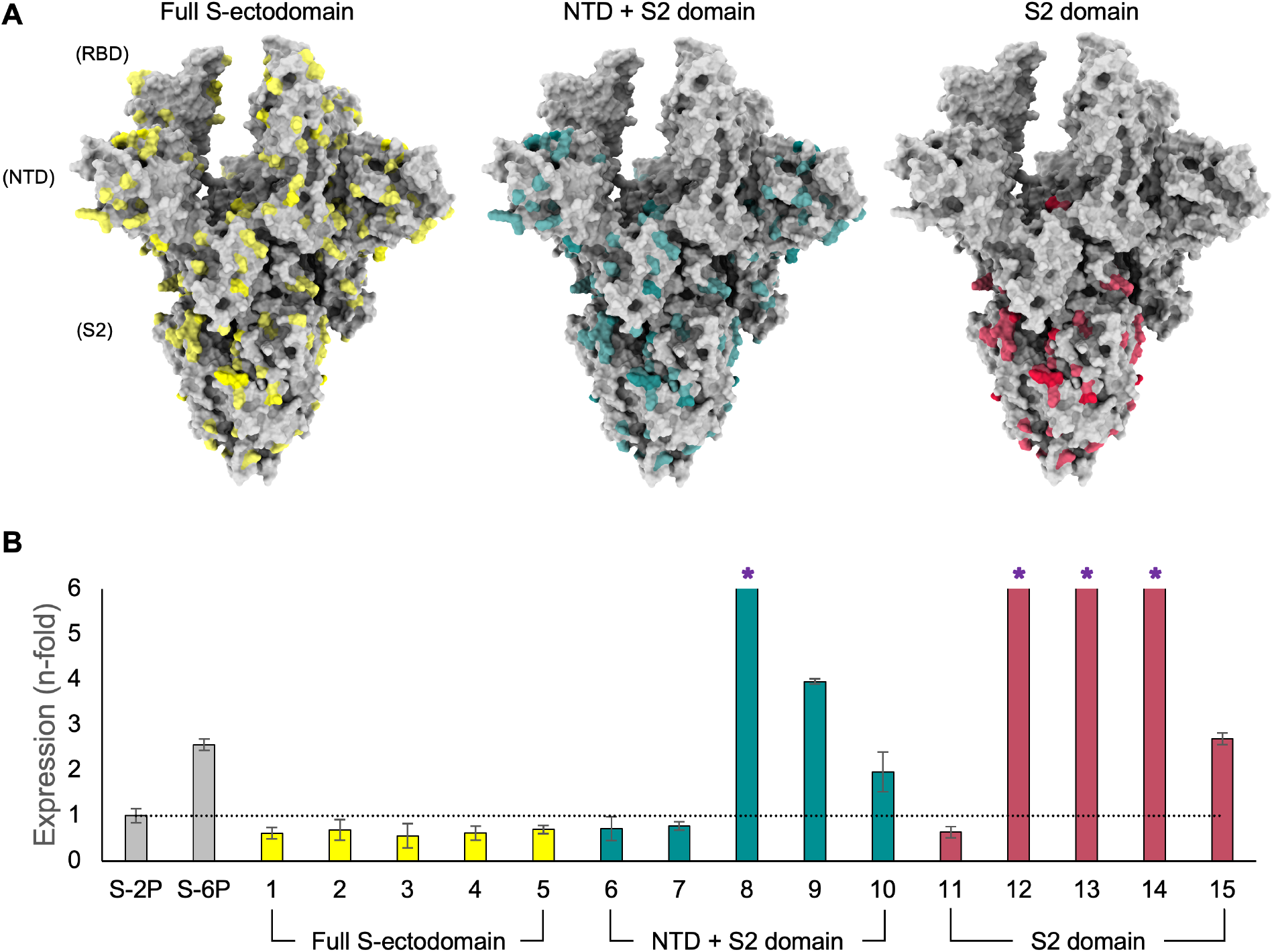
Evolutionary-based design strategy and expression levels of spike mutants. **(A)** Model of the SARS-CoV-2 Spike (S) protein structure with all three RBDs in the open conformation. The first design strategy (left) allowed for mutations (yellow) across the full S-ectodomain. The second design strategy (center) limited the design landscape to the NTD + S2 domain (mutations in teal). The third design strategy (right) only modified the S2 domain and mutations are colored red. **(B)** Protein expression level (determined by biolayer interferometry using anti-HIS biosensors) of Spike mutants normalized to S-2P (shown by dotted line) and grouped according to domain-specific design strategies in (A). S-6P (HexaPro) was used as an additional comparator. Error bars were determined from duplicate measurements and asterisks indicate designs where expression exceeded the assays limit of quantitation.

Due to the difficulty of modelling large and dynamic proteins such as S, and facing the significant sequence diversity in coronavirus families, we decided to use three independent strategies for the final design step in order to increase the probability of identifying sequences that yield desirable protein characteristics. The first strategy incorporated mutations across the full S-ectodomain, the second strategy limited the design space to the NTD + S2 domain, and the third strategy further restricted the mutations to the S2 domain (**Fig. 1A**). A total of thirty-six designs (twelve constructs per strategy) were created, where each design contained from 28 to 141 mutations (full S-ectodomain), 17 to 103 mutations (NTD + S2), or 12 to 25 mutations (S2 domain). To narrow the candidate pool for *in vitro* analysis, the twelve designs per strategy were placed into five groups based on phylogenetic analysis and we selected one representative of each group for production and purification, followed by biochemical and biophysical characterization (**Fig. S1A-C**).

### Expression, antigenicity, and stability of Spike designs

Genes containing sequences for each of the fifteen selected constructs were cloned into a mammalian expression vector and tested for expression in HEK293 cells. Among the fifteen designs selected, seven (three for the NTD + S2, and four for the S2 domain only) displayed expression levels that exceeded S-2P, with five of those designs (numbers 8, 9, 12, 13 and 14) also having expression levels higher than S-6P (**Fig. 1B**). Specifically, design 8 (NTD + S2 domain) and designs 12, 13, and 14 (S2 domain only) displayed the largest increase in expression relative to S-2P (>6 fold) or S-6P (>3 fold) and exceeded the upper limit of quantification of the assay, while design 9 (NTD + S2 domain) had an ∼4 fold and ∼1.5-fold increase compared to S-2P and S-6P, respectively. The expression levels of designs targeting the full S-ectodomain were significantly lower than detected for S-2P (∼1.5 fold) or S-6P (∼4-fold). This is consistent with observations that despite the high tolerance for variability within the RBD, many mutations can result in deleterious effects on protein expression [59]. Designs 6 and 7 (NTD + S2 domain) and design 11 (S2 domain) also had expression levels lower than S-2P. Based on these results, full S-ectodomain designs, as well as designs 6, 7, and 11, were not considered for further biochemical and biophysical characterization, leaving seven top designs for further analysis.

The majority of the most potent SARS-CoV-2 nAbs isolated to date target epitopes spanning the surface of the RBD [21, 22]. To understand whether modifications within the NTD or S2 domain altered the presentation or accessibility of key RBD epitopes, we used surface plasmon resonance (SPR) to measure binding of the RBD-targeting antibodies CR3022 (requires at least two RBDs in the open state) and S309 (recognizes open or closed RBDs), as well as binding to the hACE2 receptor (binds a single RBD in the open conformation). Designs 8, 9, 10 (NTD + S2 domain), and designs 12, 13, and 14 (S2 domain) bound CR3022, S309, and hACE2 receptor with equilibrium dissociation binding constants, (K_D_ values), in the picomolar range and with similar binding affinity as S-2P, indicating no structural disruptions to these epitopes (**Table 1, Fig. S2 and Table S1**).

**Table 1:** Thermal stability and binding affinity measurements for Spike protein designs. The T_m_ values are the average of triplicate measurements with the exception of design 10, which was limited by protein quantity. For binding affinity values, the average and standard deviation of triplicate measurements are shown with the exception of design 15, which was limited by protein quantity.

| Designed Mutant | Design Strategy | T<sub>m</sub>1 (°C) | T<sub>m</sub>2 (°C) | Binding Affinity, K<sub>D</sub> (pM) |  |  |
| --- | --- | --- | --- | --- | --- | --- |
|  |  |  |  | ACE2 | CR3022 | S309 |
| 8 | NTD + S2 | 46.38 | 76.34 | 160 ± 14 | 27 ± 4 | 19 ± 16 |
| 9 | NTD + S2 | 48.35 | 79.65 | 150 ± 5 | 370 ± 240 | 44 ± 6 |
| 10 | NTD + S2 | 46.99 | 77.44 | 44 ± 4 | 2.4 ± 3.3 | 0.16 ± 0.07 |
| 12 | S2 domain | 43.72 | 86.86 | 100 ± 2 | 9.0 ± 0.9 | 11 ± 2 |
| 13 | S2 domain | 43.75 | 77.96 | 65 ± 1 | 2.2 ± 1.1 | 20 ± 3 |
| 14 | S2 domain | 44.18 | 78.65 | 190 ± 5 | 110 ± 51 | 70 ± 14 |
| 15 | S2 domain | 44.00 | 86.54 | N/A | N/A | N/A |
| S-2P |  | 44.13 | 77.58 | 200 ± 18 | 58 ± 26 | 70 ± 0 |

To determine the impact of thermal stress on the designs, purified proteins were subjected to a differential scanning fluorimetry (DSF) assay. S-2P has previously been shown to have two distinct melting transitions (T_m_1 and T_m_2) with an increase in T_m_1 being indicative of improved stability [38]. All three NTD + S2 designs (8, 9, 10) had T_m_1 values greater than S-2P (T_m_1 = 44 °C), with design 9 having the largest, yet modest, increase of 4.2 °C (T_m_1 = 48 °C) (**Table 1**). T_m_1 values for all four S2 domain designs (12, 13, 14, and 15) were comparable to S-2P. The T_m_2 values for most designs were also comparable to S-2P, with only designs 12 and 15 showing an ∼10 °C increase in melting temperature, though the impact for this improvement is not clear. Altogether, this data shows that compared to S-2P, the leading seven designs have greater expression, are as antigenic, and have either similar or improved thermostability.

### Design 14 forms stable prefusion trimers

Binding affinity measurements confirmed that the local RBD-focused antigenicity of the seven designs remained intact. To ensure that modifications to the S antigen did not inadvertently alter S2 domain architecture or overall protein morphology, we used a Glacios 200 kV electron microscope to quickly assess one construct from the NTD + S2 designs and one construct from the S2 domain designs. Design 9 was chosen from the NTD + S2 designs based on the high expression and thermostability; however, particles resembling the prefusion S trimer were infrequently observed on the cryo-EM micrographs (**Fig. S3A**). This may be due to dissociation into monomers or other misfolded states during storage or vitrification, consistent with known issues of cold handling with Spike proteins [60]. From the S2 domain designs, we selected design 14 based on the high level of protein expression and the similar thermostability and antigenicity as S-2P. Cryo-EM micrographs of design 14 revealed the expected S protein trimers with the anticipated particle size and secondary structural features (**Fig. 2A and Fig. S3B**).

**Fig 2.**
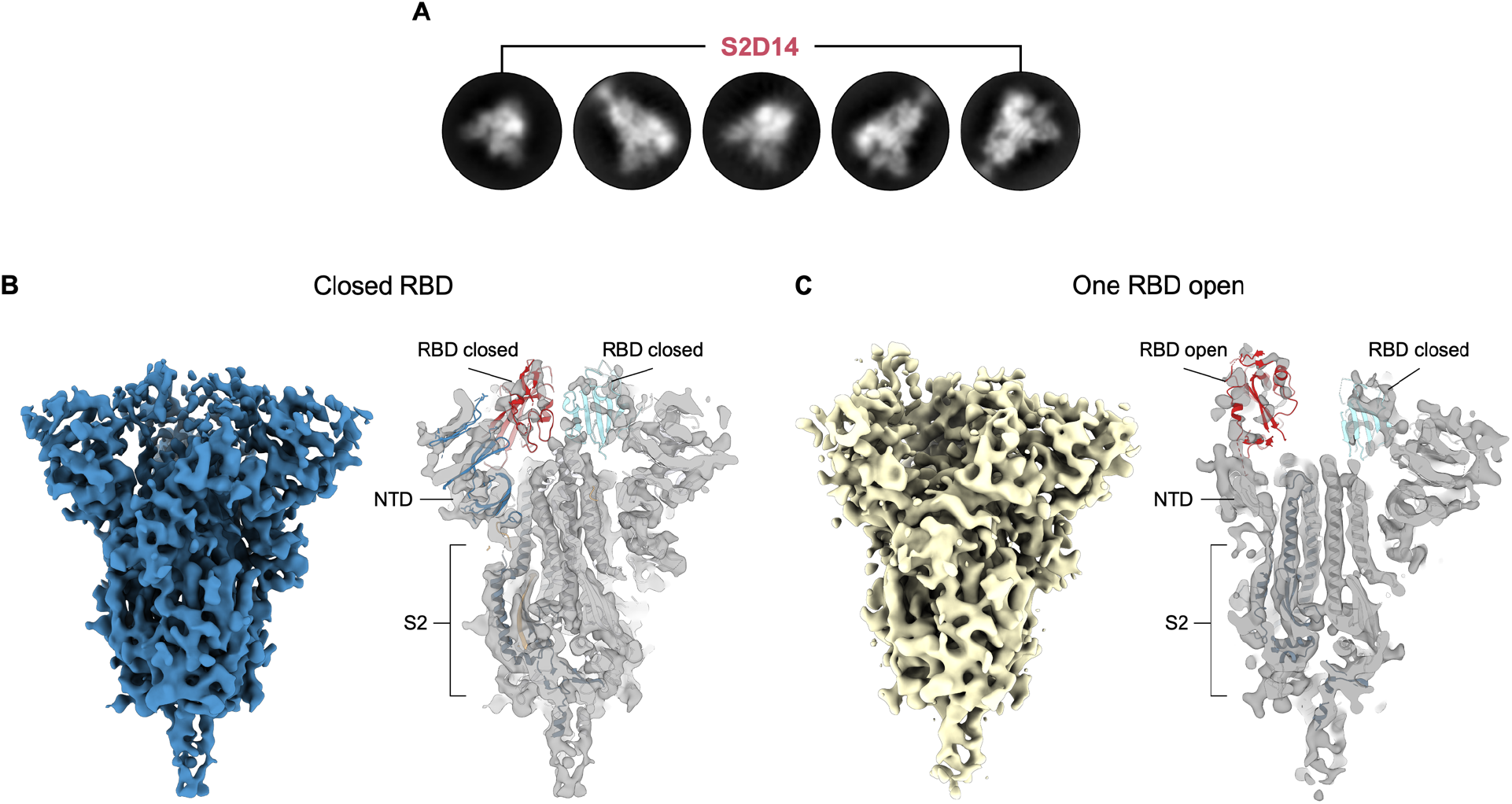
S2D14 (design 14) adopts the trimeric prefusion S conformation and exhibits RBD conformational heterogeneity. **(A)** Representative 2D classes of S2 domain design S2D14 from cryo-EM micrographs, confirming the expected S protein trimers. **(B)** Low resolution 3D reconstruction of S2D14 in a three RBD closed state (7.1 Å) is shown on the left and colored blue. Shown on the right is a cross section of the EM map with a rigid body fit of the S protein in the closed conformation (PDB 6VXX). **(C)** Low resolution 3D reconstruction of S2D14 with one RBD in the open conformation (8.5 Å) and obtained from the same dataset as (B), is shown on the left and colored yellow. Shown on the right is a cross section of the EM map with a rigid body fit of the Spike protein in a one RBD open state (PDB 6VSB).

Two distinct conformational states for design 14 were identified during 3D classification; one conformation with all three RBDs in the closed state and the second conformation with a single RBD domain in the open state (**Fig. 2B-C and Fig. S4**). Refinement of each population resulted in 7.1 Å resolution (three RBD closed) and 8.5 Å resolution (one RBD open) maps, respectively (**Table 2 and Fig. 2B-C**). The sampling of RBD open states is consistent with the recognition of the hACE2 receptor and mAb CR3022. Docking of previously published structures for either the closed Spike trimer (PDB 6VXX) or a single RBD open trimer (PDB 6VSB) into the corresponding electron density maps revealed a high degree of similarity across the S2 domains for either model, indicating that engineered S2 mutations did not disrupt the overall S protein architecture. At this point, design 14 was considered our top candidate for further evaluation and renamed S2D14 for simplicity.

**Table 2:** Cryo-EM data collection and image processing statistics for S2D14.

| Data collection |  |  |  |  |  |
| --- | --- | --- | --- | --- | --- |
| Microscope | Glacios TEM |  | Titan Krios |  |  |
| Voltage (kV) | 200 |  | 300 |  |  |
| Detector | Falcon 3 |  | Falcon 4 |  |  |
| Magnification | 120,000 |  | 120,000 |  |  |
| Pixel size (Å/pixel) | 0.91 |  | 0.67 |  |  |
| Exposure (e/Å²) | 48 |  | 43 |  |  |
| No. frames | 50 |  | 42 |  |  |
| Defocus range (µm) | 0.75 – 2.5 |  | 0.8 – 2.2 |  |  |
| Software | EPU |  | EPU |  |  |
| Total micrographs | 1,425 |  | 6,169 |  |  |
| Data processing |  |  |  |  |  |
| Particles extracted | 92,933 |  | 724,002 |  |  |
| After 2D classification | 44,459 |  | 566,825 |  |  |
|  | 3-RBD down | 1-RBD open | 3-RBD down | 2-RBD exposed | 2-RBD open |
| Particles in refinement | 7,759 | 9,154 | 33,173 | 47,773 | 87,614 |
| Symmetry | C3 | C1 | C3 | C1 | C1 |
| Final Resolution | 7.1 | 8.5 | 2.8 | 3.3 | 3.1 |
| PDB ID |  |  | 8EPN | 8EPQ | 8EPP |
| EMDB ID |  |  | 28531 | 28533 | 28532 |
| Model Statistics |  |  |  |  |  |
| MolProbity score |  |  | 1.5 | 1.60 | 1.67 |
| Clash score |  |  | 6.70 | 7.20 | 7.58 |
| Rotamer outliers (%) |  |  | 0.84 | 0.12 | 0.35 |
| Ramachandran Plot |  |  |  |  |  |
| Favored (%) |  |  | 97.34 | 96.75 | 96.21 |
| Allowed (%) |  |  | 2.66 | 3.21 | 3.65 |
| Disallowed (%) |  |  | 0.00 | 0.03 | 0.14 |

### Immunization with S2D14 elicits high neutralizing antibody titers

To assess the immunogenicity of S2D14 compared to S-2P, we immunized BALB/c mice with AS03 (oil-in-water emulsion) adjuvanted proteins at either 0.3 μg or 3.0 μg doses. Intramuscular injections were given on days zero and twenty-one, with serum collected three weeks post-I injection and two weeks post-II injection (**Fig. 3A**). Total anti-Spike IgG titers (assessed by anti-Spike ELISA) indicated that S2D14 was immunogenic at both the 0.3 μg and 3.0 μg dose and that titers were boosted after the second dose **(Fig. 3B-C)**. Anti-Spike IgG titers were comparable between S-2P and S2D14 at both post-I and post-II time points at the 0.3 μg dose, but for the 3.0 μg dose an increase in IgG titers was observed for S2D14 compared to S-2P at the post-II time point (**Fig. 3B-C**).

**Fig 3.**
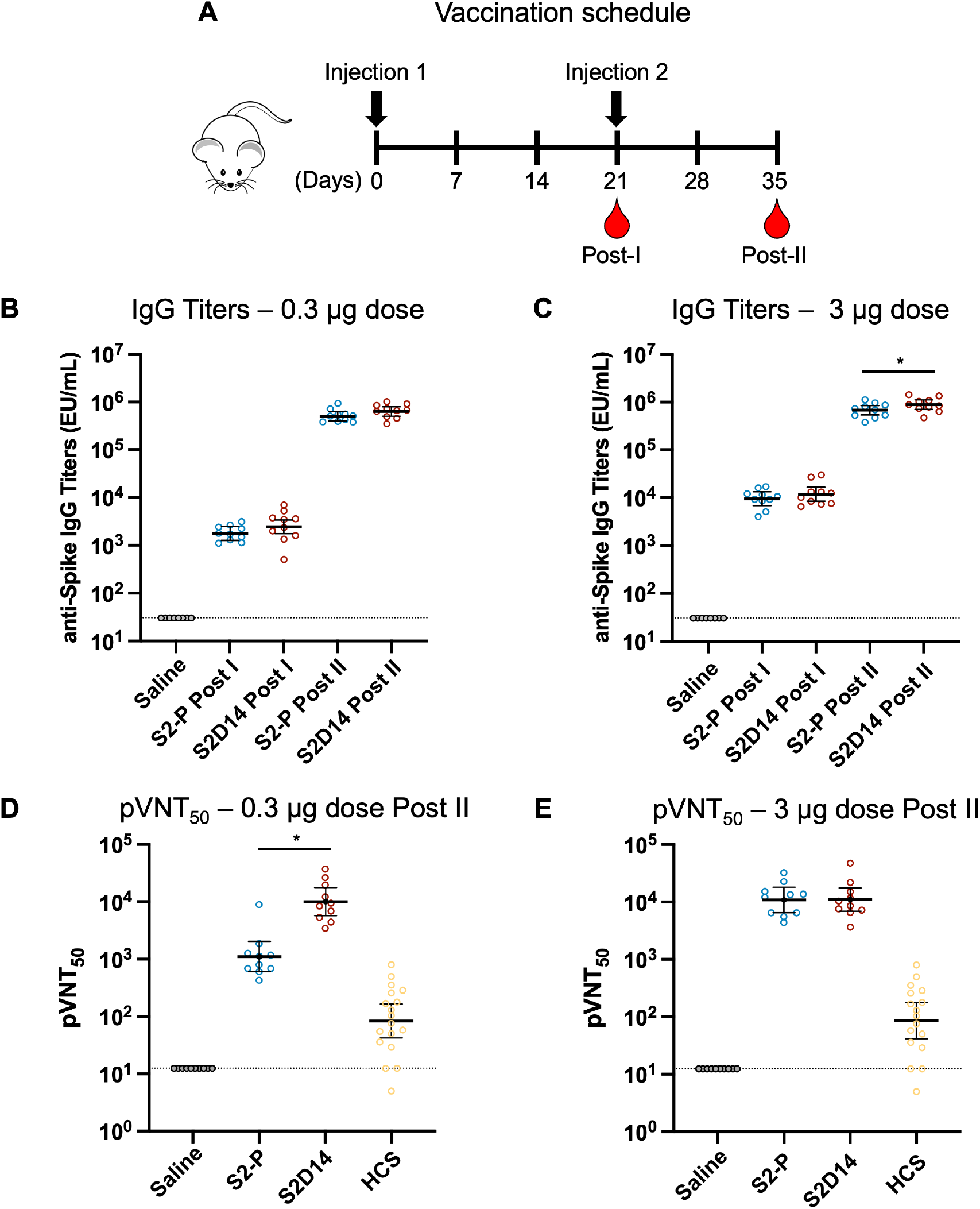
IgG and neutralizing antibody titers from mice immunized with adjuvanted S2D14 or S-2P. **(A)** Mouse immunization schedule. Mice were immunized with either a 0.3 μg or 3.0 μg dose containing AS03 adjuvanted S2D14 or S-2P. Serum was collected three weeks after the first immunization (post-I) and two weeks after the second immunization (post-II). **(B and C)** ELISA IgG titers of S2D14 compared to S-2P at both 0.3 μg **(B)** and 3.0 μg **(C)** dose for both post-I and post-II serum collections. Individual data with GMT and 95% CI are presented. **(D and E)** Neutralizing antibody titers against the Wuhan strain were analyzed 2 weeks post-II at two antigen dosages, 0.3 μg **(D)** and 3.0 μg **(E)**. Neutralizing antibody titers at either dose post-II were noticeably greater than for human convalescent sera (HCS). Individual data with GMT and 95% CI are presented. For HCS samples, GMT and 95% CI were calculated separately from vaccination groups in Prism GraphPad. Asterisks, *, indicate statistically significant increase in neutralization response based on GMR comparisons with a 2-sided 90% CI. Ratios for which the CI does not include 1 are considered statistically significant. See Methods for a description of the sera panel.

S2D14 elicited a robust neutralizing antibody response to the Wuhan strain at post-II for either vaccine dose **(Fig. 3D-E)**. Importantly, at post-II for the 0.3 μg dose, S2D14 induced statistically significant higher neutralization antibody titers compared to S-2P (9.1-fold increase). Additionally, both S-2P and S2D14 had low neutralizing titers post-I at either dose (**Fig. S5**), but noticeably higher neutralizing titers at either dose at post-II than for human convalescent serum (HCS). (**Fig. 3D-E**) [61].

Next, we determined neutralizing antibody titers against variants of concern that were available at the time of this study: Alpha (B1.1.7), Beta (B.1.351), Delta (B.1.617.2), and Omicron (BA.1) strains **(Fig. 4A-B)**. S2D14 elicited a neutralizing antibody response to all the variant strains when immunized at both the 0.3 μg and 3.0 μg doses but with an overall decrease in neutralizing antibody titers compared to Wuhan strain for Beta (∼10-fold), Delta (∼5-fold), and Omicron (∼100-fold) variants. Compared to S-2P at post-II for the 0.3 μg dose, S2D14 induced statistically significant higher neutralization antibody responses against the Alpha (10.8-fold increase), Beta (8.2-fold increase) and Delta (23.4-fold increase) variants (**Fig. 4A)**. Regarding Omicron strain, although many mice did not show measurable titers (4 out of 10 in S2D14 group and 7 out of 10 in S-2P group), and the difference was not statistically significant, at 0.3 μg dose S2D14 did elicit an ∼10-fold higher neutralizing antibody response than S-2P. At post-II for the 3.0 μg dose, S2D14 elicited statistically significant higher neutralizing antibody responses against the Alpha and Beta variants but were comparable to S-2P for Delta and Omicron variants **(Fig. 4B)**. Altogether, this data shows that S2D14 is immunogenic and capable of eliciting high levels of IgG binding antibodies (Wuhan) and neutralizing antibodies (Wuhan and VoC strains). Additionally, although the total IgG titers are similar between S2D14 and S-2P after vaccination with a 0.3 μg dose, the antibody response elicited by S2D14 at this dose is superior in neutralization potency and breadth.

**Fig 4.**
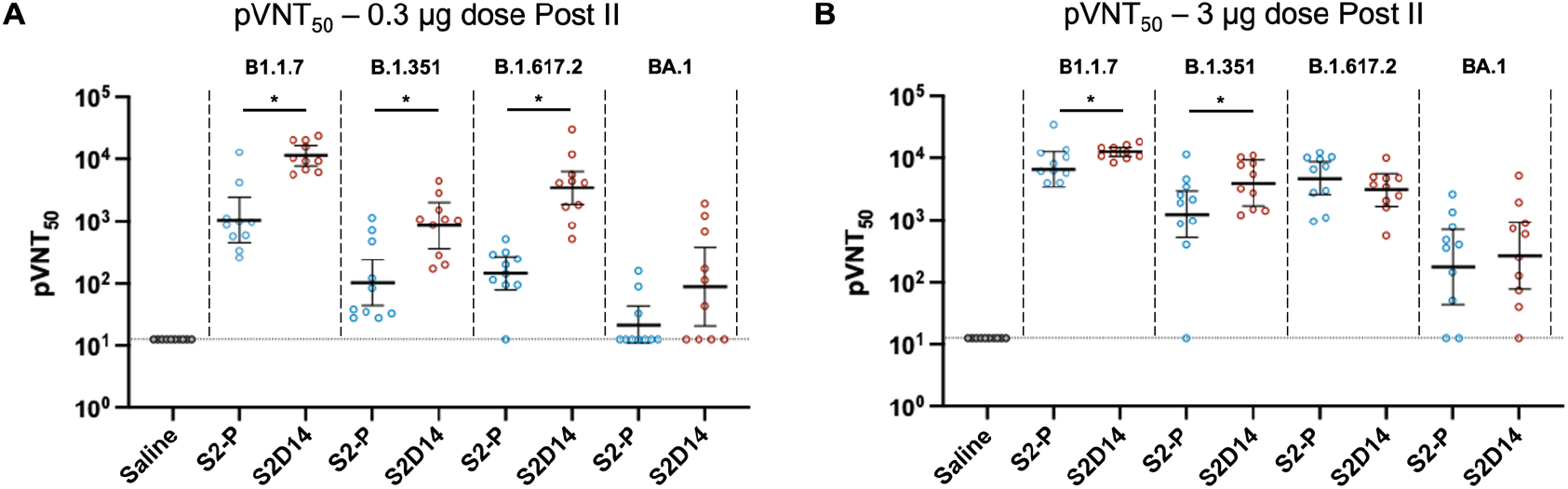
S2D14 elicits neutralizing antibody titers in mice capable of neutralizing variants of concern. **(A)** Neutralization of variants of concern compared between S2D14 and S-2P at low dose for post-II serum collection. S2D14 induced statistically significant higher neutralization responses against Alpha (B1.1.7), Beta (B.1.351) and Delta (B.1.617.2) variants compared to S-2P. **(B)** Neutralization of VoCs compared between S2D14 and S-2P at high dose for post-II serum collection. S2D14 induced statistically significant higher neutralization responses against the Alpha (B1.1.7) and Beta (B.1.351) variants compared to S-2P. Individual data with GMT and 95% CI are presented. Asterisks, *, indicate statistically significant increase in neutralization response based on GMR comparisons with a 2-sided 90% CI. Ratios for which the CI does not include 1 are considered statistically significant.

### Cryo-EM structures of S2D14 reveal conformational bias towards RBD-open states

To better understand the molecular basis for the improved breadth and immunogenicity of S2D14, a more extensive cryo-EM dataset was collected for high-resolution structural analysis using a Titan Krios 300 kV electron microscope. From a single dataset, we were able to determine the structure of S2D14 in three distinct conformations: two RBDs open (43% of particles, 3.1 Å resolution); three RBDs down (16% of total particles, 2.8 Å resolution); and two RBDs exposed with one RBD down (23% of total particles, 3.3 Å resolution) (**Fig. 5A-D, Table 2, and Fig. S6**). Although this final class most closely aligned to the one RBD open mask used during focused classification, two of the RBDs lacked clear density suggesting a state with two dynamic RBDs, each sampling an RBD open state, yet distinguishable from the two RBD open structure (**Fig. 5B-D**). Altogether, these conformations reveal that ∼66% of particles contain at least 2 RBD’s in the open or exposed states. The two RBD open conformation was not identified for S-2P; rather, a single RBD in the open conformation was reported, suggesting this to be the dominant class [20]. Docking of Fabs S309 [62], S2X259 or S2K146 [25, 26] onto S2D14 confirm that neutralization sensitive RBD epitopes, in particular those exposed only in an RBD open state (recognized by S2X259 and S2K146), are accessible with no apparent clashes (**Fig. 5E**).

**Fig. 5.**
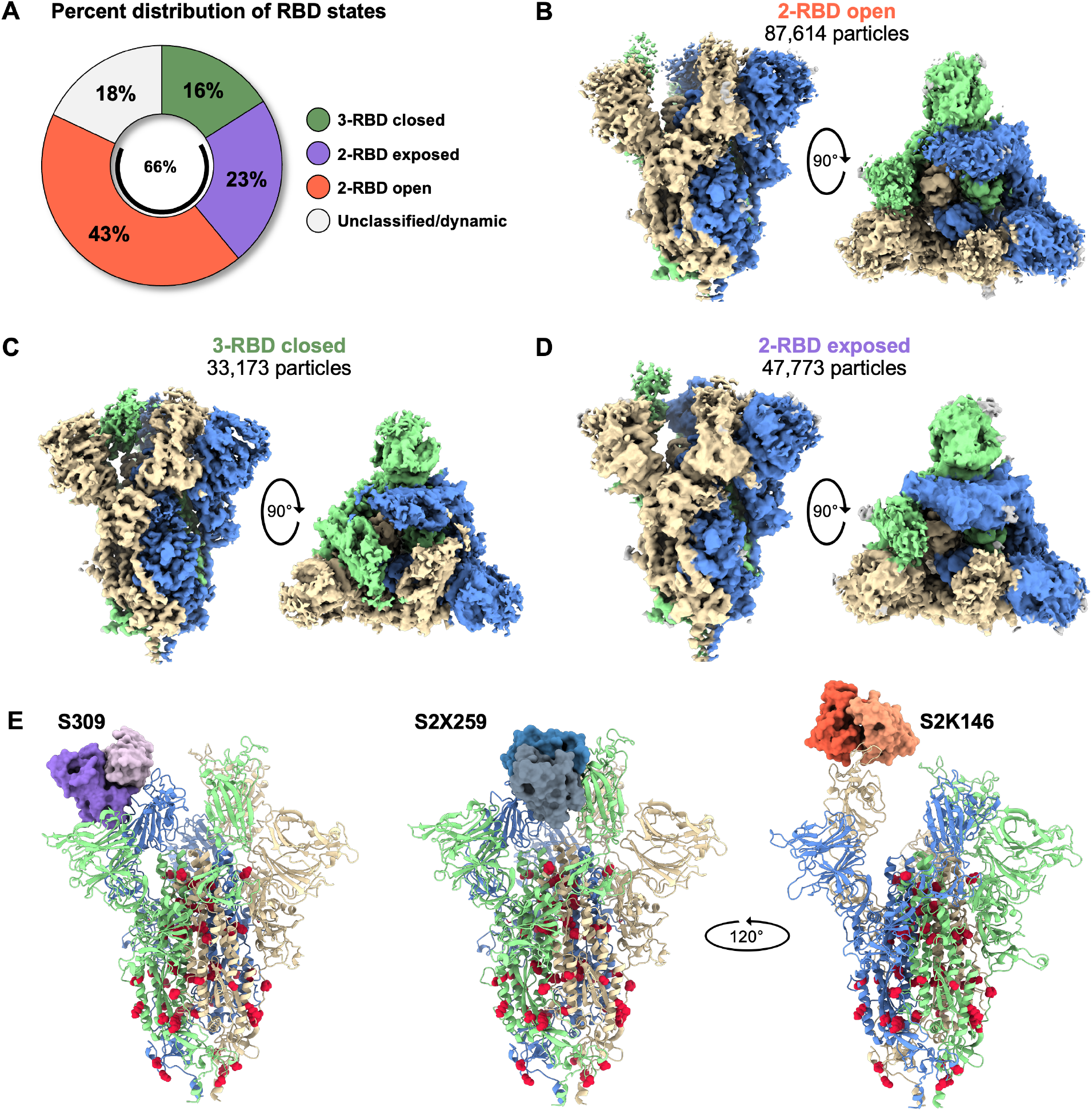
S2D14 favors RBD open conformational states and is accessible for binding RBD nAbs. **(A)** Pie chart illustrating the percent distribution of RBD conformational states. (**B)** Cryo-EM structure of S2D14 in the two RBD open conformation (3.1 Å resolution) was the dominant population and represents 43% of total particles. **(C)** Cryo-EM structure of S2D14 in the three RBD down conformation (2.8 Å resolution) was the minority population with 16% of total particles. **(D)** Cryo-EM structure (3.3 Å resolution) of S2D14 in the two RBD exposed state was representative of 23% of the total population. 18% of the particles could not be classified into a distinct conformation. Side and top views are shown for each structure, and monomers are individually colored for clarity. **(E)** Docking of Fabs of S309 (left), S2X259 (center), and S2K146 (right) onto S2D14 in the two RBD open conformation confirms the accessibility of these broadly neutralizing RBD epitopes. S2D14 protomers are colored blue, green, or tan, and mutations are shown as red spheres.

The twenty-one mutations (twenty S2 domain and D614G mutations) present in S2D14 are well resolved with clear side chain densities (**Fig. 6A-C**). Analysis of the structures provide molecular details for the three main types of mutations incorporated by evolutionary consensus design: a) improvement of complementary electrostatic potential between interprotomer contact regions (**Fig. 6A**); b) optimization of hydrophobicity and van der Waals (VDW) contacts (**Fig. 6B**); and/or c) introduction of hydrogen bond interactions (**Fig. 6C**). For example, the mutation T998N, introduces a hydrogen bond between the asparagine oxygen with side chain of Y756, located on the same protomer, while the mutation T1027E within the central helix forms a hydrogen bonding network between the E1027 side chain and R1039 of an adjacent S protomer which may help to promote stable trimer formation (**Fig. 7A-C**). Several hydrogen bonds are also lost compared to S-2P, due to mutations such as T734V, S1003A, and Q1005N (**Fig. S7A-B**). However, as shown by DSF, the overall thermostability of S2D14 is the same as S-2P, suggesting that this loss of energy is well compensated not only by the formation of new hydrogen bonds, but also by the VDWs and electrostatic mutations that are introduced. One example for the addition of an electrostatic interaction is the A701E mutation which positions the E701 side chain within ∼5 Å of the of K786 side chain of an adjacent S protomer (**Fig. 7D and Fig. S7C-E**).

**Fig 6.**
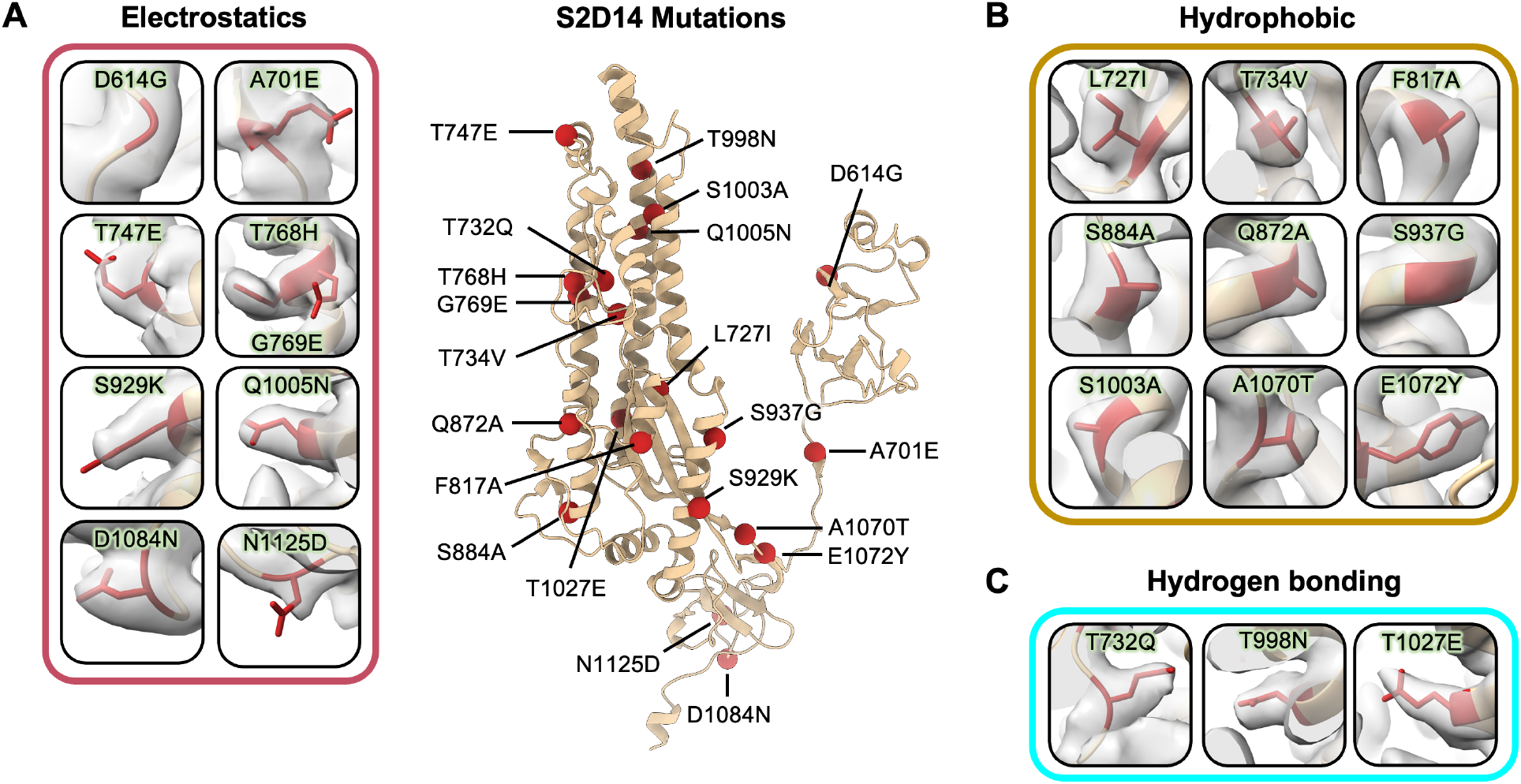
PROSS mutations in S2D14 are well resolved by cryo-EM. A single protomer of S2D14 residues 600-1147 are shown in the center in ribbon representation with mutations indicated as red spheres. Mutations are grouped based on changes in either **(A)** electrostatic complementarity, **(B)** alterations in hydrophobicity, or **(C)** introduction of hydrogen bonds. Residue side chains are depicted as sticks and cryo-EM density is shown as transparent surface.

**Fig 7.**
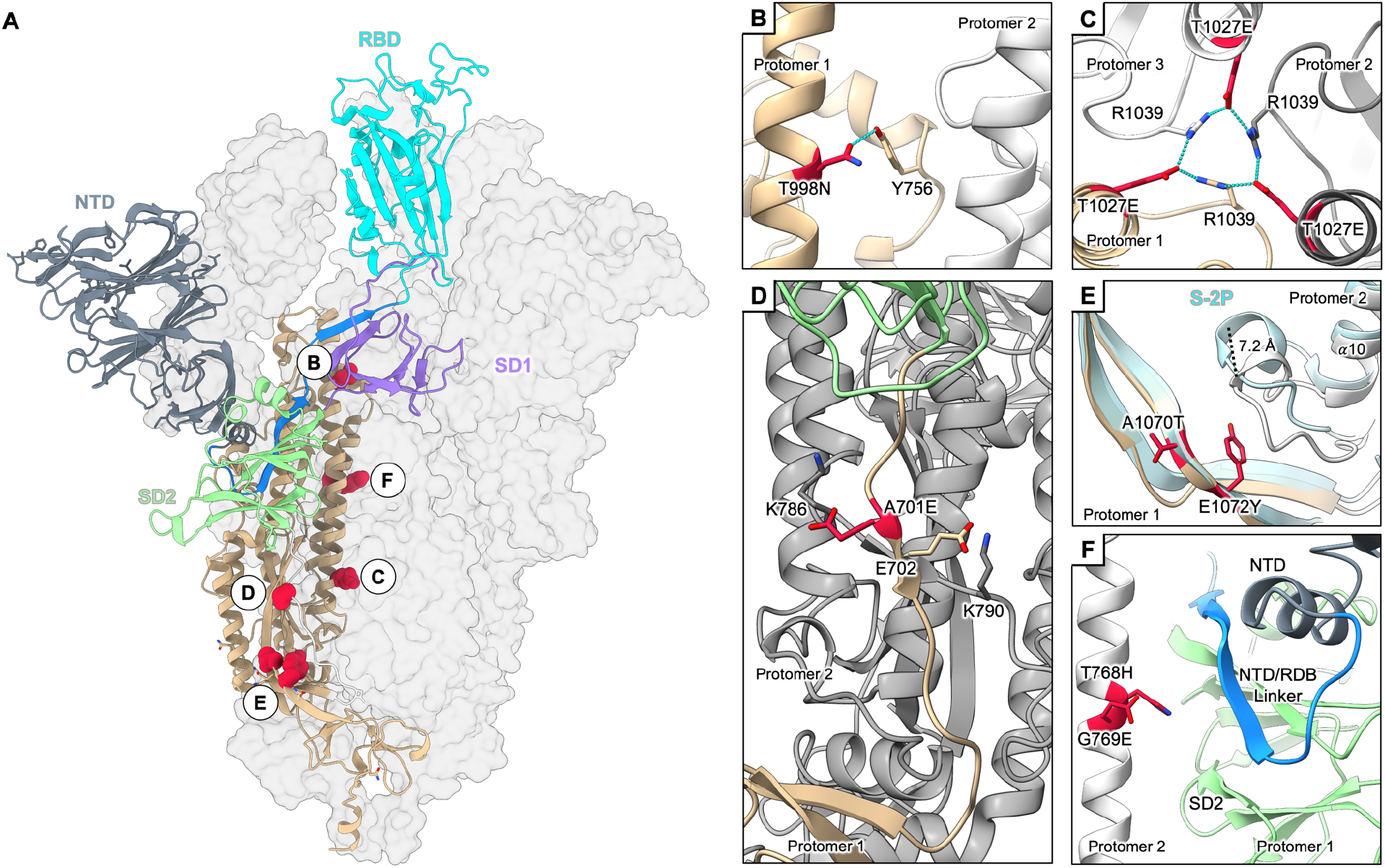
Mutations within S2D14 that may influence RBD open states. **(A)** S2D14 is shown with two protomers as transparent surfaces, and a single protomer is depicted as cartoon with domains colored as labeled in the figure. Mutations that may stabilize the RBD open conformation are colored red and shown as spheres. **(B)** A T998N mutation in the central helix domain introduces a hydrogen bond with Y756 within the same protomer. (**C)** A T1027E mutation in the S2 core forms an interprotomer hydrogen bond network between R1039 residues. (**D)** An A701E mutation creates a negative patch at the S1/S2 linker region in proximity to a positively charged lysine residue K786 on an adjacent protomer. (**E)** The E1072Y mutation creates a hydrophobic surface which displaces an adjacent loop away from the trimer interior by ∼7.2 Å. The displacement of this loop, which is adjacent to HR1, may partially destabilize the S2 domain core and influence opening of the RBD. The positioning of this loop for S-2P is shown for comparison (light blue). **(F)** T768H and G769E mutations alter the local surface potential in proximity to the NTD/RBD linker (blue) and SD2 domain (green) which acts as a hinge for RBD opening.

Several of the mutations help rationalize why S2D14 has a preference for the RBD open state. Superposition of the S2 domain (residues 821–1147) of S2D14 in the two RBD open conformation with S-2P reveals an RMSD of 1.6 Å over all C_α_ atoms, indicating that modification of the S2 domain did not dramatically alter the S2 subunit architecture (**Fig. S8**). Despite this high structural similarity, there is a marked difference in the positioning of the loop connecting the *α*10 and *α*11 helices (residues G885-Q895) near mutations A1070T and E1072Y located on the adjacent S protomer. These mutations reduce the overall charge of the interface between these two S protomers, creating a more hydrophobic patch which displaces the loop connecting *α*10 and *α*11 by 7.2 Å (measured from the C_*α*_ atom of residue A890) (**Fig. 7E and Fig. S9A-E**). Modification of this loop, particularly with a A892P mutation, has been shown to provide an increase in expression and increased conformational sampling of RBD states [38, 63]. Additionally, the mutations T768H and G769E insert charged residues which alter the surface potential of the alpha helix (residues 759-782) that packs against the SD1 domain of a second S protomer (**Fig. 7F and Fig. S9F-H**). This subdomain acts as a hinge for opening of the RBD domain; therefore, this change in surface charge may destabilize the SD1 domain, further explaining the preference for an RBD open state.

## Discussion

The emergence of SARS-CoV-2 variants has heightened the concerns about the efficacy of currently approved vaccines, which are based on the prefusion stabilized S protein antigen, S-2P [1-4, 64]. Taking a computational approach starting with S-2P modelled to have three RBDs in the open conformation, we used evolutionary consensus design for the *in silico* optimization of trimeric S antigens. These antigens were then biochemically, biophysically, structurally, and immunogenically characterized to assess the impact of mutagenesis. This strategy led to the selection of design 14, named S2D14, which has more than six-fold higher expression in HEK293 cells and maintains similar thermostability and antigenicity as S-2P. Cryo-electron microscopy structures revealed that roughly two-thirds of the S2D14 particles have either one or more RBD’s in an open or exposed state, whereas the reported S-2P cryo-EM structure has a dominant class of particles with only a single RBD in the open state [20]. The increased RBD exposure for S2D14 translated to a post vaccination immunogenic response in mice with higher levels of neutralizing antibodies against Wuhan and VoC strains.

A preference for the RBD open state has previously been reported for the S protein design u1S2q, where the authors showed that modification of interdomain contacts resulted in 67% of particles in either a one RBD or two RBD open conformation [19]. Mutations for u1S2q were focused on contact regions between subdomains and selected using a rational-based approach that resulted in four mutations between the SD1 hinge and S2 domain. Interestingly, although we observed a similar proportion of S proteins in an RBD open or exposed state (66%), none of the twenty-one mutations identified by consensus design in S2D14 overlap with mutations in u1S2q, suggesting there are multiple mechanisms for modulating RBD dynamics.

One possible mutation in S2D14 which may explain the RBD open preference is D614G, which interestingly, is now a dominant mutation within all VoCs. The mutation has been proposed to energetically favor the RBD open state [65], due to the loss of a salt bridge between D614 and K854 in the fusion peptide-proximal region. This could be why VoCs harboring this mutation are now prevailing, as D614G mediates a more efficient ACE-2 interaction and improves infectivity of the virus [66, 67]. The ability of antibodies elicited by S2D14 immunization to neutralize VoCs more efficiently than those elicited by S-2P immunization provides convincing evidence that the design of an S protein trimer with increased RBD accessible states is important for generating a more broadly protective immune response.

Currently, efforts to target Omicron variants by delivery of booster vaccines of variant-matched sequences have shown reduced immunogenicity and inferior protection against Omicron infection in non-human primates compared to vaccine boosts using the approved mRNA vaccine (Wuhan strain) [13, 68].

This highlights an urgent need for improved vaccines capable of broad protection again VoCs. Some strategies taken to improve protective efficacy have included focusing the immune response towards the RBD through multivalent display of the RBD on nanoparticles, or through the modification of the RBD sequence [27, 28]. In the case of nanoparticles, mosaic display of RBDs from diverse sarbecoviruses was capable of neutralizing SARS-CoV-2 variants, including Omicron, and was protective against challenge with SARS-2 Delta variant and SARS-1 in NHPs [27]. Alternatively, modification of the RBD sequence was shown to focus the immune response to potently neutralizing epitopes and to elicit neutralizing titers greater than the unmodified RBD in mice [28]. Aside from the RBD, vulnerable epitopes have also been described for the NTD and SD1 [21, 22, 69-77]. The S2 domain, in particular, is highly conserved among coronaviruses and it was found that non-SARS-CoV-2 exposed individuals had IgG that bound the CoV-2 S2 domain but not S1, perhaps by previous infection with a related human coronavirus [78]. Recently, stabilization of the MERS-CoV S2 domain was shown to elicit cross-reactive betacoronavirus antibodies in mice suggesting that the S2 domain has potential to elicit broad coronavirus protection [52].

Antibody responses against other vulnerable sites on S, such as the NTD, S1 subdomains, or the S2 domain are also expected to play a role in the improved neutralization potency of S2D14. Electron microscopy-based polyclonal epitope mapping, which has described the antibody landscape for other vaccine targets such as HIV-1 Envelope [79-81], and Influenza HA [82], has recently been applied to characterize antibody specificities against the Spike protein in SARS-CoV-2 convalescent sera [83]. A comparison of preexisting serum antibodies against other betacoronavirus spike proteins revealed epitopes that were differentially targeted by SARS-CoV-2 infected individuals. Similarly, a detailed molecular map of the immune response elicited by S2D14 could provide a valuable comparison between polyclonal specificities generated after vaccination or by natural infection, which in turn could facilitate future rational designs. The role of T cell immunity is another important consideration for optimized S antigens which is suggested to play a role in limiting severe-to-critical COVID-19 [84]. S-6P was shown to induce higher frequencies of antigen-specific CD8^+^ T cells producing Th1 cytokines than S-2P while trimeric RBD linked to the HR1 and HR2 domain of S2 induced RBD-specific IL-4 and INF-γ producing memory T cells [85, 86]. Future studies should aim to examine whether mutations in S2D14 might have introduced new epitopes and assess the impact of T cell immunity on limiting disease severity after immunization with S2D14.

A significant finding in our work is that although vaccination of mice with either S2D14 or S-2P results in the elicitation of similar IgG titers, S2D14 elicits a higher functional response that was more effective than S-2P in neutralizing the ancestral Wuhan strain and variants of concern (Alpha, Beta, and Delta). A recent report of the immunogenicity of recombinant VSV vectored S-6P found that the enhanced stability and expression of S-6P led to elicitation of a more effective nAb response at lower doses than S-2P against the Wuhan strain and VoC including Alpha (B.1.1.7), Gamma (P.1), Beta (B.1.351), Epsilon (B.1.427), and Delta (B.1.617.2) strains [85]. Combined with enhanced expression, vaccination with S2D14 could translate well into dose-sparing effects when delivered in a recombinant setting, but also may translate to a lower dose when delivered by alternative platforms such as mRNA or replication-competent viral vectors. Regarding neutralization of Omicron strain, immunization of mice with the 0.3 μg dose of S2D14 resulted in reduced neutralizing antibodies when compared to titers observed against the Wuhan strain but were ∼10-fold greater than titers elicited by S-2P. Therefore, S2D14 could serve as a scaffold for additional modifications which further improve nAb responses against Omicron or future variants. Overall, this work highlights the benefits of using an evolutionary consensus approach for antigen design and supports additional investigation into S2D14 as an immunogen for combating future coronaviruses.

## Methods

### Computational design

Rosetta comparative modeling (RosettaCM) [87] was used to build a model, with cyclic symmetry constraints [56], of the SARS CoV-2 S antigen with the receptor binding domain (RBD) in the open conformation (PDB Accession Numbers: 6VSB, 6VYB). The model reconstruction strategy used a combination of x-ray and cryo-EM structures (PDB Accession Numbers: 6VYB, 6VW1, 6NB7 (SARS-CoV-1)). Symmetric interface design was performed on the lowest energy RosettaCM trimer, focusing on a monomeric chain with two virtual and identical partners, adapting protocols from the Protein Repair One-Stop Shop (PROSS), with the updated beta energy scoring function [50, 51].

To design mutations of the spike protein from SARS-CoV-2 using evolutionary constraints for the introduction of stabilizing residues, homologous sequences were obtained from the non-redundant BLAST database [55] and narrowed to 500 glycoprotein sequences from various coronavirus lineages (**Data S1**). These aligned sequences were calculated into a position-specific scoring matrix (PSSM) with the PSI-BLAST algorithm [55]. The matrix represents the likelihood for each of the 20 amino acids being present at each residue position, within the aligned sequences.

The Rosetta FilterScan mover [88] was used to perform single point mutagenesis of all the residues to the preferred PSSM mutations, targeting the full spike ectodomain (S), N-terminal domain plus S2 domain (NTD + S2), or the S2 domain only. The mutation scan was binned within twelve different energy thresholds (−0.5, -1, -1.5, -2, -2.5, -3, -3.5, -4, -4.5, -5, -5.5, -6 kcal/mol) to increase mutation sequence diversity. A RosettaScripts [89] algorithm that energetically combined the proposed single mutations was used to reduce the search space, yielding twelve total stabilizing designs for each round of mutations, and representing each energy threshold.

### S-2P and Spike mutant expression and quantification

The gene encoding the sequence of designed S mutants were synthesized and cloned into an in-house mammalian expression vector pBW, with a hexahistidine sequence added to the C terminus. The Spike designs were expressed in HEK293 cells using 3.0 mL of Expi293 expression in each well of a 24 deep well high-throughput expression system. Each design was tested in duplicate wells. Cell culture supernatants were harvested on Day 5-6 when viability was approximately 50%.

The Octet quantification assay for protein expression level was performed on an Octet 96 Red system (Sartorius). Harvested cell media were centrifuged and cell supernatants were prepared in a 96-well plate with S-2P or S-6P standards diluted in media from 20 μg/mL to 0.3125 μg/mL. The standard and mutant binding curves were measured using anti-HIS biosensors, where the concentration of each mutant in media was calculated by fitting the measured initial binding rate to the calibration curve. The expression levels were measured in duplicate wells of each mutant’s media and the average readout was reported.

### Purification of mutants

The culture supernatant of selected SARS-CoV-2 S mutants produced in 1 L scale was directly loaded through 5 ml Ni-NTA Excel column (Cytiva Life Sciences). The column was washed by 300 mM sodium chloride, 50 mM imidazole in 20 mM HEPES buffer, pH 7.5 and captured S mutants were eluted by 300 mM sodium chloride, 250 mM imidazole in 20 mM HEPS buffer, pH 7.5. The collected samples from appropriate elution fractions were pooled and concentrated prior to further purification of the S trimer by gel filtration using a Superose 6 increase 10/300 GL column (Cytiva Life Sciences) with 1x PBS, pH 7.4 as running buffer. The fractions corresponding to the targeted S mutants based on SDS-PAGE analysis were pooled together and quantitated using the absorbance at 280 nm.

### Surface plasmon resonance

Surface plasmon resonance experiments were performed in a running buffer composed of 0.01 M HEPES, pH 7.4, 0.15 M NaCl, 3 mM EDTA, 0.005% v/v Surfactant P20 at 25 °C using the Biacore 8K (GE Healthcare), with a series S protein A sensor chip (GE Healthcare). The ACE2 receptor or SARS-CoV-2 spike-specific antibodies (CR3022 or S309) were immobilized on the protein A sensor chip (GE Healthcare) at a ligand capture level of ∼100 RU. Serial dilutions of purified spike designs were injected, at concentrations ranging from 10 nM to 1.25 nM. The resulting data were fit to a 1:1 binding model using the Biacore Evaluation Software (GE Healthcare).

### Differential scanning fluorimetry

Nano differential scanning fluorimetry was used to assess the thermal stability of purified spike designs on a Prometheus NT.48 instrument (NanoTemper Technologies). Samples were diluted to 0.2 mg/mL with PBS and 20 μL of each sample was loaded into capillary tubes. A temperature ramp was set to 1ºC/minute, ranging from 20 ºC to 95 ºC. The reported values are the mean of the 1^st^ derivative of the ratio of intrinsic tryptophan fluorescence emission wavelengths for protein unfolding/folding (350 nm/330 nm), measured in triplicate.

### Mouse immunization studies

An *in vivo* study was performed to assess the immunogenicity of S2D14 compared to S-2P in a mouse model. Female BALB/c mice, 7-8 weeks of age at the start of the study, were immunized (N=10 mice/group) with AS03 (an oil-in-water emulsion adjuvant system containing 1.186 mg alpha-tocopherol/dose)-adjuvanted Spike proteins at two dosage levels of 3.0 μg or 0.3 μg [90]. A saline placebo control group was also included in the study (N=4) but was not considered for statistical analysis. Spike proteins and AS03 were admixed shortly before injection. Mice were injected intramuscularly twice 3-weeks apart and bled 3 weeks after the initial immunization (post-I) and 2 weeks after the second immunization (post-II).

The *in vivo* study was conducted in accordance with the GSK Policy on the Care, Welfare and Treatment of Laboratory Animals and were reviewed by the Institutional Animal Care and Use Committee by the ethical review process at the institution where the work was performed. All studies followed ARRIVE Guidelines as applicable and were conducted in compliance with provisions of the USDA Animal Welfare Act, the Public Health Service Policy on Humane Care and Use of Laboratory Animals and the U.S. Interagency Research Animal Committee Principles for the Utilization and Care of Research Animals.

### Neutralization assays and ELISA

The serum CoV2-specific antibody responses were assessed using a Wuhan strain pseudo-virus neutralization assay to measure functional antibodies as described previously [61] and an ELISA (pre-fusion S-2P antigen absorbed to the solid phase) to measure IgG binding antibodies for all mice (Nexelis, Laval, Quebec). Human SARS-CoV-2 infection/COVID-19 convalescent sera (n=22) were obtained at CHU Tivoli, Belgium, from donors 23-61 years of age, mostly from females, after PCR-confirmed diagnosis and at least 28 days after the participants were asymptomatic. Human samples were obtained with informed consent. All recruitment, sample collection, and experimental procedures using human samples have been approved by relevant institutional review boards and by GSK human sample management board. Neutralizing antibodies in post-II serum samples from mice immunized with S-2P, and S2D14 or placebo were additionally measured using a pseudo-virus neutralization assay with the Alpha, Beta, Delta, and Omicron variant strains (Nexelis, Laval, Quebec).

### Statistical analysis of IgG binding and neutralization data

To assess differences between S-2P and S2D14 vaccines, for IgG binding titers, an ANOVA model for repeated measures was fitted on log10-transformed data including vaccine, dose, time, and their interaction as fixed factors and considering homogeneity of variances between groups. For post-II neutralization data, except with Omicron strain, an ANOVA model was fitting on log10-transformed data with vaccine and dose as fixed factors. For the Omicron assay, a model accounting for left censored data was fitted on log10-transformed data. Homogeneity of variances between study groups was considered for all except Alpha and Omicron assays.

Geometric means of titers (GMT) with corresponding 95% confidence intervals as well as geometric mean ratios (GMR) and 90% confidence intervals (2-sided test with alpha=0.05) were computed from these models to compare responses to S-2P and S2D14 vaccines by dose. GMRs for which the confidence interval doesn’t include 1 are considered statistically significant. The analyses included more vaccination groups than reported (14 for IgG data and 6 for neutralization data) as the *in vivo* study initially consisted of additional vaccinated groups irrelevant to this study, but multiplicity of comparisons was not considered. GMT with corresponding 95% confidence intervals for HCS samples were computed separately from vaccination groups using Prism (GraphPad Software, San Diego, California USA).

### Cryo-EM sample preparation

SARS-CoV-2 S designs 9 and S2D14, previously isolated by nickel-affinity chromatography, were further purified by SEC using a Superose 6 10/300 GL column (Cytiva Life Sciences) with TBS buffer composed of 10 mM Tris pH 7.5, 150 mM NaCl. Fractions containing purified spike protein were diluted to a final concentration of 0.4 mg/mL prior to specimen preparation on EM grids. 3.0 μL of prepared samples were then applied to glow discharged Quantifoil 1.2/1.3 - 400 mesh copper grids, blotted, and plunged into liquid ethane using an FEI Vitrobot Mark IV vitrification apparatus set at 100% relative humidity and 4 °C (ThermoFisher Scientific).

### Cryo-EM image processing and modeling

#### Glacios TEM

A total of 1,425 movies were collected using a FEI Glacios TEM at 200 kV equipped with a Falcon 3 direct electron detector at a magnification of 120,000 corresponding to a pixel size of 0.91 Å/pixel. A more detailed description of imaging parameter used during data collection can be found in Table 2.

Cryo-EM single particle analysis was carried out using the RELION 3.1 image processing suite [91]. Briefly, frame alignment was performed using RELION’s implementation of MotionCor2 followed by CTF estimation using CTFFIND4.1 [92, 93]. A subset of images was used to manually select particles to build a set of 2D templates for automated particle picking across the entire dataset. Using the 2D templates for auto-picking, a total of 92,933 particles were selected from the full set of images. Particles were next subjected to 2D classification, which resulted in the selection of 44,459 particles. Next, 3D classification was performed against a reference structure built using RELION’s *de novo* 3D model generation and applying C3 symmetry. Three classes retained the trimeric morphology, and the fourth class was discarded due to poor structure quality. The three classes consisting of 39,718 particles were subjected to an additional round of 3D classification using C1 symmetry, which resulted in one class of particles with three RBDs in a closed state (7,759 particles) and a second class with one RBD in the open state (9,154 particles).

For particles in the closed conformation, 3D refinement was performed using C3 symmetry, while C1 symmetry was used for 3D refinement for the one RBD open conformation. After initial 3D refinement, both datasets were followed by CTF-refinement to correct for beam aberrations and determine per-particle defocus values. Bayesian polishing of the CTF-refined particles was then performed to correct for per-particle motion. A final 3D refinement of both structures converged to 7.1 Å for the RBD closed conformation and 8.5 Å for the single RBD open conformation according to the gold-standard 0.143 FSC criteria.

#### Titan Krios

A total of 6,169 movies were collected using an FEI Titan Krios operating at 300 kV and equipped with a Falcon 4 direct electron detector (Nanosciences Center, Cambridge University, UK) at a magnification of 120,000 corresponding to a pixel size of 0.67 Å/pixel. A more detailed description of settings used for imaging during data collection can be found in Table 2.

Cryo-EM single particle analysis was carried out using the RELION 3.1 image processing suite [91]. For a visual depiction of the single particle workflow, refer to Fig. S6. Briefly, alignment of the raw movie frames was performed using RELION’s implementation of MotionCor2 followed by CTF estimation using CTFFIND4.1 [92, 93]. Next, a subset of particles was manually selected and used to generate 2D classes that were then used as templates for particle picking across the entire set of 6,169 micrographs, which resulted in the selection of 724,002 particles. Particles were initially extracted at 4X binning with a pixel size of 2.68 Å/pixel. The binned particle stack was subjected to a single round of 2D classification and classes showing features consistent with the S trimer were selected for further processing. The 566,825 particles selected from 2D classification were subjected to one round of 3D classification against a reference model that was generated using RELION’s *de novo* 3D model generation. Among the generated classes was a single population resembling the known trimeric S morphology. This stack of 205,674 particles was then re-extracted as the unbinned particle set. A consensus C1 refinement was performed to estimate correct particle poses resulting in a 3.8 Å resolution cryo-EM map. Focused 3D classification was then performed using individual masks encompassing the NTD and RBD for RBDs in the closed state, one RBD open, and two RBD open conformation, which resulted in separation of the particles into three distinct classes: three RBD closed class (33,173 particles), two RBD exposed class (47,773 particles) and a two RBD open class (87,614 particles). Initial 3D auto-refinement for the three RBD closed, two RBD exposed, and two RBD open conformations resulted in 3.9 Å, 3.5 Å and 3.5 Å resolution cryo-EM maps respectively. 3D-refined particles for each set were further subjected to CTF refinement and Bayesian polishing to estimate per-particle defocus values and correct for individual particle motions. A final 3D refinement resulted in an improved resolution for each map according to the gold-standard FSC 0.143 criteria. Final resolutions for the three RBD down, two RBD exposed, and two RBD open conformations were 2.8 Å, 3.3 Å, and 3.1 Å respectively.

Modeling of each cryo-EM structure was performed using Phenix and COOT [94, 95]. For the three RBD down conformation, PDB 6VXX was used as a starting structure where mutations for S2D14 were inserted using ChimeraX [17, 96]. For the RBD-exposed and RBD-open structures, a model for the two RBD open conformation was built from PDB 6VSB and mutations for S2D14 were inserted using ChimeraX [20]. For each model and corresponding map, real space refinement was performed in Phenix, initially by applying morphing, simulated annealing and applying secondary structure restraints. To rebuild loops missing in the original PDB files and fit into the corresponding cryo-EM density, RosettaCM was performed using the initial Phenix models and structural templates for known NTD and RBD structures (PDB 7LY3 and 7DEU). Once the full homology model was built, Rosetta Cartesian refinement was performed [97], followed by torsional refinement to build glycan structures [98, 99]. A final real space refinement applying minimization, ADP refinement and occupancy optimization was performed using Phenix. Loops, disordered regions, and glycans lacking structural information in the cryo-EM density were removed in the resulting Rosetta models.

## Supplemental online material

Supplemental figures S1-S9 and tables S1.

## Acknowledgments

The authors would like to thank Karen Matsuoka for expression, harvesting and buffer exchange of culture supernatants, Nurjahan Mehzabeen and Claire Harelson for the purification of Spike antigens, and Laëtitia Leloup for her assistance with statistical analysis of the antibody data. Pascal Cadot coordinated the manuscript development

## Data availability

Data supporting the findings of this manuscript are available from the corresponding authors upon reasonable request. Cryo-EM maps of S2D14 in the two RBD open, RBD exposed and RBD closed conformations have been deposited in the Electron Microscopy Data Bank as EMD-28532, EMD-28533, and EMDB-28531 respectively. The atomic coordinates for the Cryo-EM structures of S2D14 in the two RBD open, two RBD exposed and RBD closed conformations have been deposited to the RCSB Protein Data Bank, PDB 8EPP, PDB 8EPQ, and PDB 8EPN, respectively.

## Author contributions

WH, NW, MB, EM, MJB, JW, AMS, CM, GM, YH, CC, LL were involved in the conception and design of the study and/or the development of the study protocol. JW, ST, LL, AB, LC, CM participated to the acquisition of data. WH, NW, MB, JW, ST, LL, CC, CM, GM, and SV analyzed and interpreted the results. All authors were involved in drafting the manuscript or revising it critically for important intellectual content. All authors had full access to the data and approved the manuscript before it was submitted by the corresponding author.

## References

1. Volz, E., et al., Assessing transmissibility of SARS-CoV-2 lineage B.1.1.7 in England. Nature, 2021. 593(7858): p. 266–269.

2. Tegally, H., et al., Detection of a SARS-CoV-2 variant of concern in South Africa. Nature, 2021. 592(7854): p. 438–443.

3. Jhun, H., et al., SARS-CoV-2 Delta (B.1.617.2) Variant: A Unique T478K Mutation in Receptor Binding Motif (RBM) of Spike Gene. Immune Netw, 2021. 21(5): p. e32.

4. Zhou, H., et al., Resistance of SARS-CoV-2 Omicron BA.1 and BA.2 Variants to Vaccine-Elicited Sera and Therapeutic Monoclonal Antibodies. Viruses, 2022. 14(6).

5. Jangra, S., et al., SARS-CoV-2 spike E484K mutation reduces antibody neutralisation. Lancet Microbe, 2021. 2(7): p. e283–e284.

6. Pouwels, K.B., et al., Effect of Delta variant on viral burden and vaccine effectiveness against new SARS-CoV-2 infections in the UK. Nat Med, 2021. 27(12): p. 2127–2135.

7. Cao, Y., et al., BA.2.12.1, BA.4 and BA.5 escape antibodies elicited by Omicron infection. Nature, 2022. 608(7923): p. 593–602.

8. Cao, Y., et al., Imprinted SARS-CoV-2 humoral immunity induces convergent Omicron RBD evolution. bioRxiv, 2022.

9. Wang, X., et al., Homologous or heterologous booster of inactivated vaccine reduces SARS-CoV-2 Omicron variant escape from neutralizing antibodies. Emerg Microbes Infect, 2022. 11(1): p. 477–481.

10. Zhang, Z., et al., A Heterologous V-01 or Variant-Matched Bivalent V-01D-351 Booster following Primary Series of Inactivated Vaccine Enhances the Neutralizing Capacity against SARS-CoV-2 Delta and Omicron Strains. J Clin Med, 2022. 11(14).

11. Scheaffer, S.M., et al., Bivalent SARS-CoV-2 mRNA vaccines increase breadth of neutralization and protect against the BA.5 Omicron variant. bioRxiv, 2022.

12. Muik, A., et al., Exposure to BA.4/BA.5 Spike glycoprotein drives pan-Omicron neutralization in vaccine-experienced humans and mice. bioRxiv, 2022.

13. Tubiana, J., et al., Reduced antigenicity of Omicron lowers host serologic response. bioRxiv, 2022.

14. Andreano, E. and R. Rappuoli, SARS-CoV-2 escaped natural immunity, raising questions about vaccines and therapies. Nat Med, 2021. 27(5): p. 759–761.

15. Cao, Y., et al., Omicron escapes the majority of existing SARS-CoV-2 neutralizing antibodies. Nature, 2022. 602(7898): p. 657–663.

16. Bosch, B.J., et al., The coronavirus spike protein is a class I virus fusion protein: structural and functional characterization of the fusion core complex. J Virol, 2003. 77(16): p. 8801–11.

17. Walls, A.C., et al., Structure, Function, and Antigenicity of the SARS-CoV-2 Spike Glycoprotein. Cell, 2020. 183(6): p. 1735.

18. Benton, D.J., et al., Receptor binding and priming of the spike protein of SARS-CoV-2 for membrane fusion. Nature, 2020. 588(7837): p. 327–330.

19. Henderson, R., et al., Controlling the SARS-CoV-2 spike glycoprotein conformation. Nat Struct Mol Biol, 2020. 27(10): p. 925–933.

20. Wrapp, D., et al., Cryo-EM structure of the 2019-nCoV spike in the prefusion conformation. Science, 2020. 367(6483): p. 1260–1263.

21. Starr, T.N., et al., SARS-CoV-2 RBD antibodies that maximize breadth and resistance to escape. Nature, 2021. 597(7874): p. 97–102.

22. Premkumar, L., et al., The receptor binding domain of the viral spike protein is an immunodominant and highly specific target of antibodies in SARS-CoV-2 patients. Sci Immunol, 2020. 5(48).

23. Andreano, E., et al., Hybrid immunity improves B cells and antibodies against SARS-CoV-2 variants. Nature, 2021. 600(7889): p. 530–535.

24. Andreano, E., et al., Extremely potent human monoclonal antibodies from COVID-19 convalescent patients. Cell, 2021. 184(7): p. 1821–1835 e16.

25. Tortorici, M.A., et al., Broad sarbecovirus neutralization by a human monoclonal antibody. Nature, 2021. 597(7874): p. 103–108.

26. Park, Y.J., et al., Antibody-mediated broad sarbecovirus neutralization through ACE2 molecular mimicry. Science, 2022. 375(6579): p. 449–454.

27. Cohen, A.A., et al., Mosaic RBD nanoparticles protect against challenge by diverse sarbecoviruses in animal models. Science, 2022. 377(6606): p. eabq0839.

28. Dickey, T.H., et al., Design of the SARS-CoV-2 RBD vaccine antigen improves neutralizing antibody response. Sci Adv, 2022. 8(37): p. eabq8276.

29. Sun, W., et al., The self-assembled nanoparticle-based trimeric RBD mRNA vaccine elicits robust and durable protective immunity against SARS-CoV-2 in mice. Signal Transduct Target Ther, 2021. 6(1): p. 340.

30. Walls, A.C., et al., Elicitation of Potent Neutralizing Antibody Responses by Designed Protein Nanoparticle Vaccines for SARS-CoV-2. Cell, 2020. 183(5): p. 1367–1382 e17.

31. Dai, L., et al., Efficacy and Safety of the RBD-Dimer-Based Covid-19 Vaccine ZF2001 in Adults. N Engl J Med, 2022. 386(22): p. 2097–2111.

32. Kaabi, N.A., et al., Safety and immunogenicity of a hybrid-type vaccine booster in BBIBP-CorV recipients in a randomized phase 2 trial. Nat Commun, 2022. 13(1): p. 3654.

33. Kaabi, N.A., et al., Immunogenicity and safety of NVSI-06-07 as a heterologous booster after priming with BBIBP-CorV: a phase 2 trial. Signal Transduct Target Ther, 2022. 7(1): p. 172.

34. Zhang, J., et al., A mosaic-type trimeric RBD-based COVID-19 vaccine candidate induces potent neutralization against Omicron and other SARS-CoV-2 variants. Elife, 2022. 11.

35. Hernandez-Bernal, F., et al., Safety, tolerability, and immunogenicity of a SARS-CoV-2 recombinant spike RBD protein vaccine: A randomised, double-blind, placebo-controlled, phase 1-2 clinical trial (ABDALA Study). EClinicalMedicine, 2022. 46: p. 101383.

36. Eugenia-Toledo-Romani, M., et al., Safety and immunogenicity of anti-SARS CoV-2 vaccine SOBERANA 02 in homologous or heterologous scheme: Open label phase I and phase IIa clinical trials. Vaccine, 2022. 40(31): p. 4220–4230.

37. Arunachalam, P.S., et al., Adjuvanting a subunit COVID-19 vaccine to induce protective immunity. Nature, 2021. 594(7862): p. 253–258.

38. Hsieh, C.L., et al., Structure-based design of prefusion-stabilized SARS-CoV-2 spikes. Science, 2020. 369(6510): p. 1501–1505.

39. McLellan, J.S., et al., Structure-based design of a fusion glycoprotein vaccine for respiratory syncytial virus. Science, 2013. 342(6158): p. 592–8.

40. Battles, M.B., et al., Structure and immunogenicity of pre-fusion-stabilized human metapneumovirus F glycoprotein. Nat Commun, 2017. 8(1): p. 1528.

41. Hsieh, C.L., et al., Structure-based design of prefusion-stabilized human metapneumovirus fusion proteins. Nat Commun, 2022. 13(1): p. 1299.

42. Stewart-Jones, G.B.E., et al., Interprotomer disulfide-stabilized variants of the human metapneumovirus fusion glycoprotein induce high titer-neutralizing responses. Proc Natl Acad Sci U S A, 2021. 118(39).

43. Godley, L., et al., Introduction of intersubunit disulfide bonds in the membrane-distal region of the influenza hemagglutinin abolishes membrane fusion activity. Cell, 1992. 68(4): p. 635–45.

44. Sanders, R.W., et al., Stabilization of the soluble, cleaved, trimeric form of the envelope glycoprotein complex of human immunodeficiency virus type 1. J Virol, 2002. 76(17): p. 8875–89.

45. Goldenzweig, A., et al., Automated Structure- and Sequence-Based Design of Proteins for High Bacterial Expression and Stability. Mol Cell, 2016. 63(2): p. 337–346.

46. Frappier, V. and A.E. Keating, Data-driven computational protein design. Curr Opin Struct Biol, 2021. 69: p. 63–69.

47. Marks, D.S., et al., Protein 3D structure computed from evolutionary sequence variation. PLoS One, 2011. 6(12): p. e28766.

48. Hopf, T.A., et al., Mutation effects predicted from sequence co-variation. Nat Biotechnol, 2017. 35(2): p. 128–135.

49. Riesselman, A.J., J.B. Ingraham, and D.S. Marks, Deep generative models of genetic variation capture the effects of mutations. Nat Methods, 2018. 15(10): p. 816–822.

50. Park, H., et al., Simultaneous Optimization of Biomolecular Energy Functions on Features from Small Molecules and Macromolecules. J Chem Theory Comput, 2016. 12(12): p. 6201–6212.

51. Alford, R.F., et al., The Rosetta All-Atom Energy Function for Macromolecular Modeling and Design. J Chem Theory Comput, 2017. 13(6): p. 3031–3048.

52. Hsieh, C.L., et al., Stabilized coronavirus spike stem elicits a broadly protective antibody. Cell Rep, 2021. 37(5): p. 109929.

53. Lapidoth, G., et al., Highly active enzymes by automated combinatorial backbone assembly and sequence design. Nat Commun, 2018. 9(1): p. 2780.

54. Malladi, S.K., et al., One-step sequence and structure-guided optimization of HIV-1 envelope gp140. Curr Res Struct Biol, 2020. 2: p. 45–55.

55. Altschul, S.F., et al., Gapped BLAST and PSI-BLAST: a new generation of protein database search programs. Nucleic Acids Res, 1997. 25(17): p. 3389–402.

56. DiMaio, F., et al., Modeling symmetric macromolecular structures in Rosetta3. PLoS One, 2011. 6(6): p. e20450.

57. Korber, B., et al., Spike mutation pipeline reveals the emergence of a more transmissible form of SARS-CoV-2. bioRxiv, 2020.

58. Brufsky, A., Distinct viral clades of SARS-CoV-2: Implications for modeling of viral spread. J Med Virol, 2020. 92(9): p. 1386–1390.

59. Starr, T.N., et al., Deep Mutational Scanning of SARS-CoV-2 Receptor Binding Domain Reveals Constraints on Folding and ACE2 Binding. Cell, 2020. 182(5): p. 1295–1310 e20.

60. Edwards, R.J., et al., Cold sensitivity of the SARS-CoV-2 spike ectodomain. Nat Struct Mol Biol, 2021. 28(2): p. 128–131.

61. Maruggi, G., et al., A self-amplifying mRNA SARS-CoV-2 vaccine candidate induces safe and robust protective immunity in preclinical models. Mol Ther, 2022. 30(5): p. 1897–1912.

62. Pinto, D., et al., Cross-neutralization of SARS-CoV-2 by a human monoclonal SARS-CoV antibody. Nature, 2020. 583(7815): p. 290–295.

63. Juraszek, J., et al., Stabilizing the closed SARS-CoV-2 spike trimer. Nat Commun, 2021. 12(1): p. 244.

64. Huang, C., et al., Clinical features of patients infected with 2019 novel coronavirus in Wuhan, China. Lancet, 2020. 395(10223): p. 497–506.

65. Liu, Y., J. Liu, and P.Y. Shi, SARS-CoV-2 variants and vaccination. Zoonoses (Burlingt), 2022. 2(1).

66. Zhang, J., et al., Structural impact on SARS-CoV-2 spike protein by D614G substitution. Science, 2021. 372(6541): p. 525–530.

67. Benton, D.J., et al., The effect of the D614G substitution on the structure of the spike glycoprotein of SARS-CoV-2. Proc Natl Acad Sci U S A, 2021. 118(9).

68. Gagne, M., et al., mRNA-1273 or mRNA-Omicron boost in vaccinated macaques elicits similar B cell expansion, neutralizing responses, and protection from Omicron. Cell, 2022. 185(9): p. 1556–1571 e18.

69. Ju, B., et al., Human neutralizing antibodies elicited by SARS-CoV-2 infection. Nature, 2020. 584(7819): p. 115–119.

70. Tortorici, M.A., et al., Ultrapotent human antibodies protect against SARS-CoV-2 challenge via multiple mechanisms. Science, 2020. 370(6519): p. 950–957.

71. Byrnes, J.R., et al., Competitive SARS-CoV-2 Serology Reveals Most Antibodies Targeting the Spike Receptor-Binding Domain Compete for ACE2 Binding. mSphere, 2020. 5(5).

72. Tanaka, S., et al., Rapid identification of neutralizing antibodies against SARS-CoV-2 variants by mRNA display. Cell Rep, 2022. 38(6): p. 110348.

73. Xu, S., et al., Mapping cross-variant neutralizing sites on the SARS-CoV-2 spike protein. Emerg Microbes Infect, 2022. 11(1): p. 351–367.

74. Cerutti, G., et al., Potent SARS-CoV-2 neutralizing antibodies directed against spike N-terminal domain target a single supersite. Cell Host Microbe, 2021. 29(5): p. 819–833 e7.

75. Wang, Z., et al., Conserved Neutralizing Epitopes on the N-Terminal Domain of Variant SARS-CoV-2 Spike Proteins. bioRxiv, 2022.

76. Cerutti, G., et al., Neutralizing antibody 5-7 defines a distinct site of vulnerability in SARS-CoV-2 spike N-terminal domain. Cell Rep, 2021. 37(5): p. 109928.

77. Seow, J., et al., A neutralizing epitope on the SD1 domain of SARS-CoV-2 spike targeted following infection and vaccination. Cell Rep, 2022. 40(8): p. 111276.

78. Nguyen-Contant, P., et al., S Protein-Reactive IgG and Memory B Cell Production after Human SARS-CoV-2 Infection Includes Broad Reactivity to the S2 Subunit. mBio, 2020. 11(5).

79. Bianchi, M., et al., Electron-Microscopy-Based Epitope Mapping Defines Specificities of Polyclonal Antibodies Elicited during HIV-1 BG505 Envelope Trimer Immunization. Immunity, 2018. 49(2): p. 288–300 e8.

80. Nogal, B., et al., Mapping Polyclonal Antibody Responses in Non-human Primates Vaccinated with HIV Env Trimer Subunit Vaccines. Cell Rep, 2020. 30(11): p. 3755–3765 e7.

81. Antanasijevic, A., et al., Polyclonal antibody responses to HIV Env immunogens resolved using cryoEM. Nat Commun, 2021. 12(1): p. 4817.

82. Han, J., et al., Polyclonal epitope mapping reveals temporal dynamics and diversity of human antibody responses to H5N1 vaccination. Cell Rep, 2021. 34(4): p. 108682.

83. Bangaru, S., et al., Structural mapping of antibody landscapes to human betacoronavirus spike proteins. Sci Adv, 2022. 8(18): p. eabn2911.

84. Moss, P., The T cell immune response against SARS-CoV-2. Nat Immunol, 2022. 23(2): p. 186–193.

85. Lu, M., et al., SARS-CoV-2 prefusion spike protein stabilized by six rather than two prolines is more potent for inducing antibodies that neutralize viral variants of concern. Proc Natl Acad Sci U S A, 2022. 119(35): p. e2110105119.

86. He, C., et al., A self-assembled trimeric protein vaccine induces protective immunity against Omicron variant. Nat Commun, 2022. 13(1): p. 5459.

87. Song, Y., et al., High-resolution comparative modeling with RosettaCM. Structure, 2013. 21(10): p. 1735–42.

88. Whitehead, T.A., et al., Optimization of affinity, specificity and function of designed influenza inhibitors using deep sequencing. Nat Biotechnol, 2012. 30(6): p. 543–8.

89. Fleishman, S.J., et al., RosettaScripts: a scripting language interface to the Rosetta macromolecular modeling suite. PLoS One, 2011. 6(6): p. e20161.

90. Garcon, N., D.W. Vaughn, and A.M. Didierlaurent, Development and evaluation of AS03, an Adjuvant System containing alpha-tocopherol and squalene in an oil-in-water emulsion. Expert Rev Vaccines, 2012. 11(3): p. 349–66.

91. Scheres, S.H., RELION: implementation of a Bayesian approach to cryo-EM structure determination. J Struct Biol, 2012. 180(3): p. 519–30.

92. Zheng, S.Q., et al., MotionCor2: anisotropic correction of beam-induced motion for improved cryo-electron microscopy. Nat Methods, 2017. 14(4): p. 331–332.

93. Rohou, A. and N. Grigorieff, CTFFIND4: Fast and accurate defocus estimation from electron micrographs. J Struct Biol, 2015. 192(2): p. 216–21.

94. Liebschner, D., et al., Macromolecular structure determination using X-rays, neutrons and electrons: recent developments in Phenix. Acta Crystallogr D Struct Biol, 2019. 75(Pt 10): p. 861–877.

95. Emsley, P., et al., Features and development of Coot. Acta Crystallogr D Biol Crystallogr, 2010. 66(Pt 4): p. 486–501.

96. Pettersen, E.F., et al., UCSF ChimeraX: Structure visualization for researchers, educators, and developers. Protein Sci, 2021. 30(1): p. 70–82.

97. Conway, P., et al., Relaxation of backbone bond geometry improves protein energy landscape modeling. Protein Sci, 2014. 23(1): p. 47–55.

98. Frenz, B., et al., Automatically Fixing Errors in Glycoprotein Structures with Rosetta. Structure, 2019. 27(1): p. 134–139 e3.

99. Adolf-Bryfogle, J., et al., Growing Glycans in Rosetta: Accurate <em>de novo</em> glycan modeling, density fitting, and rational sequon design. bioRxiv, 2021.

