## Supplementary Table S1 and Figures S1 - S9 for "Structural and Computational Design of a SARS-2 Spike Antigen with Increased Receptor Binding Domain Exposure and Improved Immunogenicity"

Science Advances

AAAS

**This PDF file includes:**

Figs. S1 to S9
Tables S1

A

| Full S-Ectodomain | | |
|---|---|---|
| **CoV S Mutant** | **Selected Design** | **Rosetta score (kcal/mol)** |
| Target sequence | | -11615.951 |
| PreS_S_0_5 | | -12405.366 |
| PreS_S_1 | 1 | -12339.751 |
| PreS_S_1_5 | | -12294.3 |
| PreS_S_2 | 2 | -12315.109 |
| PreS_S_2_5 | | -12278.535 |
| PreS_S_3 | | -12251.713 |
| PreS_S_3_5 | 3 | -12213.033 |
| PreS_S_4 | | -12192.985 |
| PreS_S_4_5 | | -12159.976 |
| PreS_S_5 | 4 | -12080.037 |
| PreS_S_5_5 | | -12056.763 |
| PreS_S_6 | 5 | -12064.728 |

**Full S-Ectodomain**

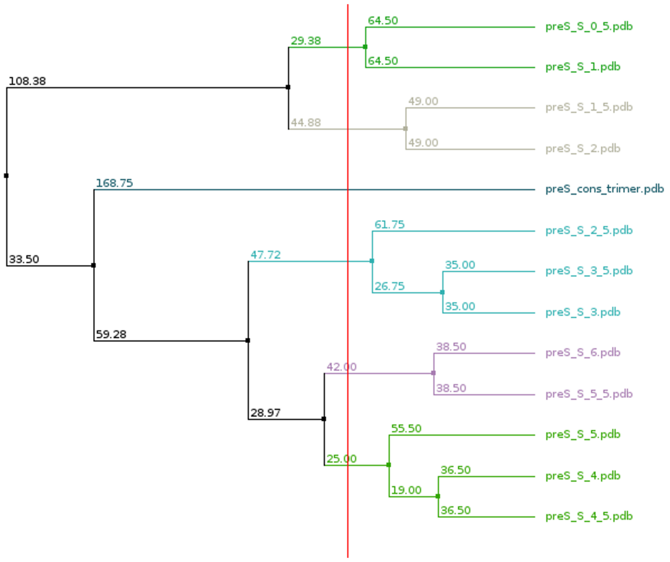


B

| NTD + S2 | | |
|---|---|---|
| **CoV S Mutant** | **Selected Design** | **Rosetta score (kcal/mol)** |
| Target sequence | | -11615.951 |
| PreS_S2_NTD_0_5 | 6 | -12142.196 |
| PreS_S2_NTD_1 | | -12202.403 |
| PreS_S2_NTD_1_5 | | -12213.696 |
| PreS_S2_NTD_2 | 7 | -12192.697 |
| PreS_S2_NTD_2_5 | | -12159.516 |
| PreS_S2_NTD_3 | 8 | -12139.76 |
| PreS_S2_NTD_3_5 | | -12078.789 |
| PreS_S2_NTD_4 | | -12064.461 |
| PreS_S2_NTD_4_5 | | -12065.383 |
| PreS_S2_NTD_5 | 9 | -12028.133 |
| PreS_S2_NTD_5_5 | | -11998.548 |
| PreS_S2_NTD_6 | 10 | -11957.319 |

**NTD + S2 domain**

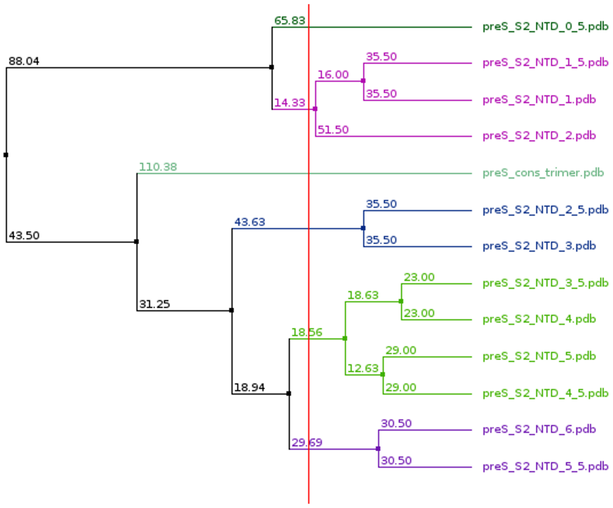

**C**

| S2 Domain | | |
|---|---|---|
| **CoV S Mutant** | **Selected Design** | **Rosetta score (kcal/mol)** |
| Target sequence | | -11615.951 |
| PreS_S2_0_5 | | -12072.375 |
| PreS_S2_1 | 11 | -12028.056 |
| PreS_S2_1_5 | | -12031.222 |
| PreS_S2_2 | 12 | -12040.859 |
| PreS_S2_2_5 | | -12035.513 |
| PreS_S2_3 | 13 | -11985.224 |
| PreS_S2_3_5 | | -11988.799 |
| PreS_S2_4 | 14 | -11969.559 |
| PreS_S2_4_5 | | -11922.035 |
| PreS_S2_5 | | -11963.851 |
| PreS_S2_5_5 | | -11928.956 |
| PreS_S2_6 | 15 | -11909.424 |

**S2 domain**

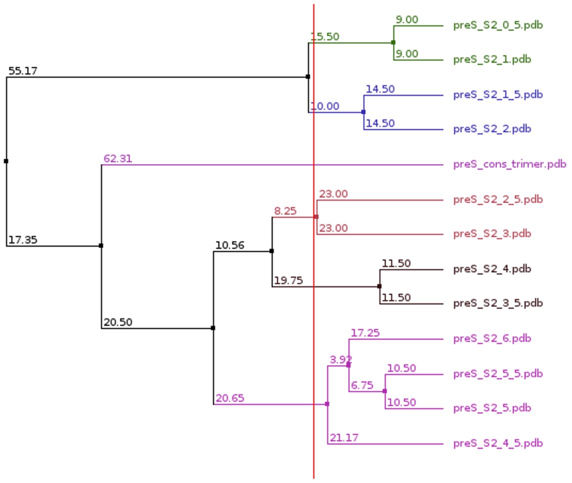


**Fig. S1: Rosetta energy scoring and selection of Spike protein designs.** Phylogenetics using BLOSUM62 sequence analysis matrix in JalView [1] and the average distance algorithm was used for each domain-specific design strategy to separate sequences into five groups (indicated by color and excluding the preS cons trimer). Since members of each group were highly similar, a single representative from each group was randomly selected, resulting in a total of 15 sequences for analysis. **(A)** Rosetta energy scores for the full S-ectodomain designs and selection of five designs, 1-5, using phylogenetics (right). **(B)** Rosetta energy scores for NTD + S2 domain designs and selection of five designs, 6-10, using phylogenetics (right). **(C)** Rosetta energy scores for S2 domain designs and selection of five designs, 11-15 using phylogenetics (right). Rosetta energy calculations for all designs were predicted to be more stable than the prefusion Spike target sequence. *preS_cons trimer (target sequence) is the S model built from multiple experimental structures and containing S-2P di-proline (K986P and V987P), and the D614G drift mutations.

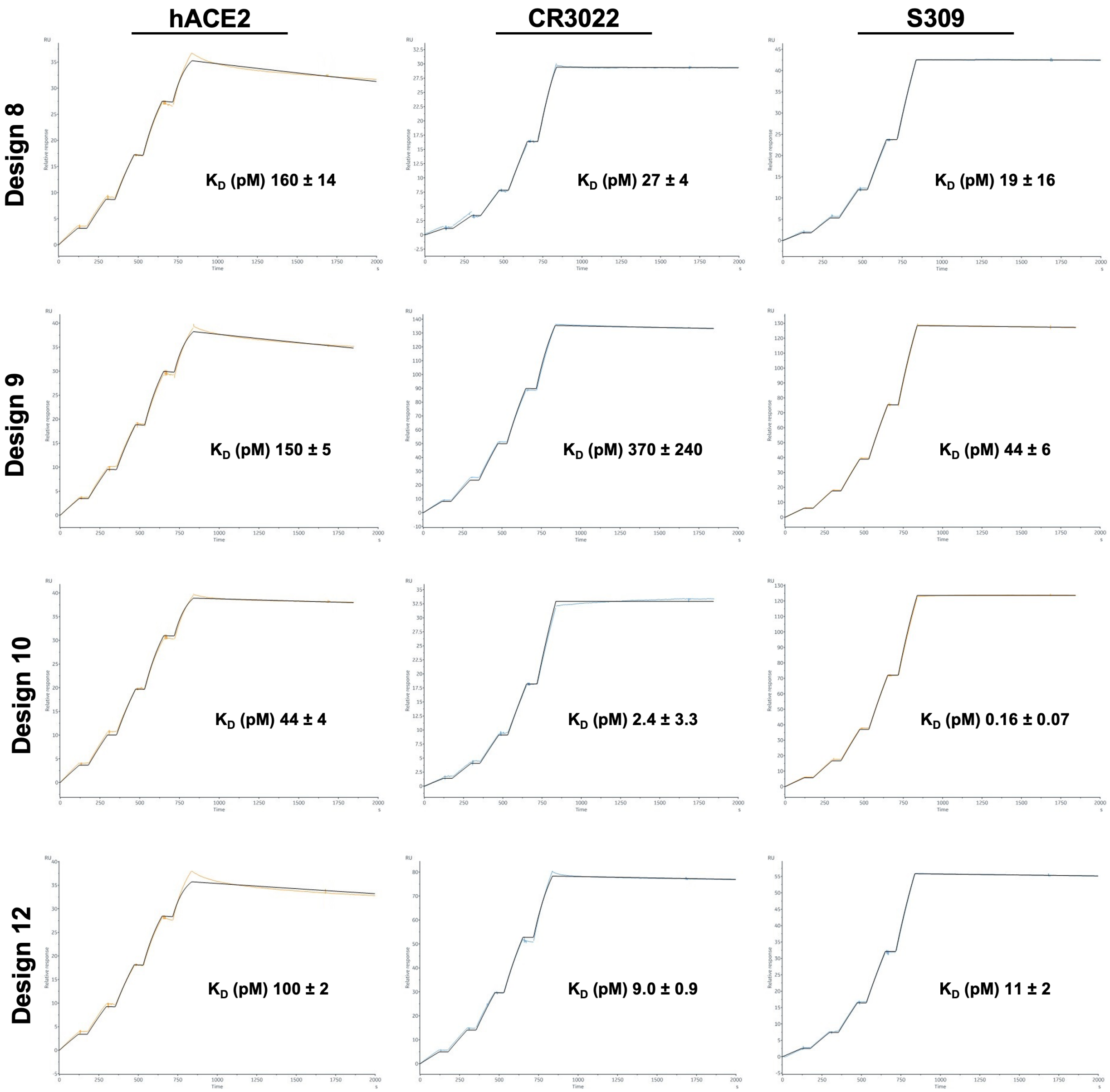
hACE2
CR3022
S309
Design 8
Design 9
Design 10
Design 12
$K_D$ (pM) 160 ± 14
$K_D$ (pM) 27 ± 4
$K_D$ (pM) 19 ± 16
$K_D$ (pM) 150 ± 5
$K_D$ (pM) 370 ± 240
$K_D$ (pM) 44 ± 6
$K_D$ (pM) 44 ± 4
$K_D$ (pM) 2.4 ± 3.3
$K_D$ (pM) 0.16 ± 0.07
$K_D$ (pM) 100 ± 2
$K_D$ (pM) 9.0 ± 0.9
$K_D$ (pM) 11 ± 2
RU
Relative response
Time
s

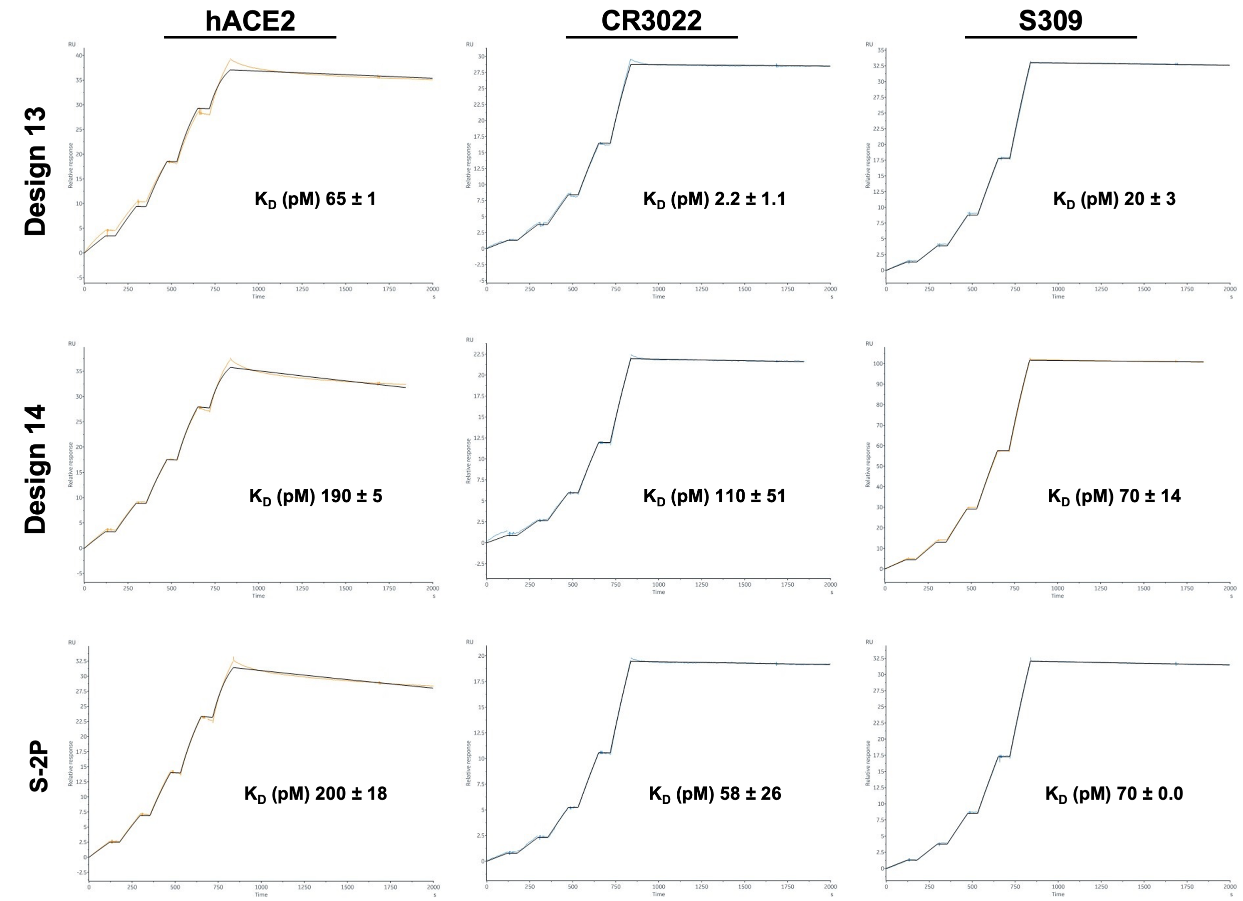


**Fig. S2: Selected SARS-CoV-2 S designs and S-2P are capable of binding hACE2 receptor, CR3022, and S309.** SPR binding curves for selected S designs and S-2P. Data were fit to a 1:1 binding model (calculated fit is shown as black line) and the average $K_D$ and standard deviations for triplicate measurements are shown below each binding curve. See Supplementary Table 1 for kinetic parameters.

| Antigen | Receptor/Binding antibody | $k_a$ (1/Ms) | $k_d$ (1/s) |
|---|---|---|---|
| S-2P | ACE2 | $4.9 \times 10^{+5} \pm 2.5 \times 10^{+4}$ | $1.0 \times 10^{-4} \pm 5.3 \times 10^{-6}$ |
| | CR3022 | $2.3 \times 10^{+5} \pm 7.4 \times 10^{+4}$ | $1.2 \times 10^{-5} \pm 2.9 \times 10^{-6}$ |
| | S309 | $2.0 \times 10^{+5} \pm 1.5 \times 10^{+4}$ | $1.4 \times 10^{-5} \pm 1.2 \times 10^{-6}$ |
| 8 | ACE2 | $5.9 \times 10^{+5} \pm 2.7 \times 10^{+4}$ | $1.0 \times 10^{-4} \pm 4.1 \times 10^{-6}$ |
| | CR3022 | $2.3 \times 10^{+5} \pm 3.1 \times 10^{+4}$ | $6.5 \times 10^{-6} \pm 1.2 \times 10^{-9}$ |
| | S309 | $2.7 \times 10^{+5} \pm 3.9 \times 10^{+4}$ | $5.6 \times 10^{-6} \pm 5.1 \times 10^{-6}$ |
| 9 | ACE2 | $5.7 \times 10^{+5} \pm 2.3 \times 10^{+4}$ | $8.9 \times 10^{-5} \pm 2.9 \times 10^{-6}$ |
| | CR3022 | $1.5 \times 10^{+5} \pm 6.4 \times 10^{+3}$ | $5.5 \times 10^{-5} \pm 3.4 \times 10^{-5}$ |
| | S309 | $1.8 \times 10^{+5} \pm 3.0 \times 10^{+3}$ | $8.0 \times 10^{-6} \pm 1.1 \times 10^{-6}$ |
| 10 | ACE2 | $6.2 \times 10^{+5} \pm 1.3 \times 10^{+3}$ | $2.7 \times 10^{-5} \pm 2.6 \times 10^{-6}$ |
| | CR3022 | $3.4 \times 10^{+5} \pm 1.7 \times 10^{+5}$ | $1.1 \times 10^{-6} \pm 1.5 \times 10^{-6}$ |
| | S309 | $1.8 \times 10^{+5} \pm 6.5 \times 10^{+3}$ | $3.0 \times 10^{-8} \pm 1.4 \times 10^{-8}$ |
| 12 | ACE2 | $6.2 \times 10^{+5} \pm 3.6 \times 10^{+3}$ | $6.3 \times 10^{-5} \pm 8.7 \times 10^{-7}$ |
| | CR3022 | $1.7 \times 10^{+6} \pm 2.3 \times 10^{+4}$ | $1.5 \times 10^{-5} \pm 1.3 \times 10^{-6}$ |
| | S309 | $8.4 \times 10^{+5} \pm 6.8 \times 10^{+4}$ | $9.4 \times 10^{-6} \pm 7.1 \times 10^{-7}$ |
| 13 | ACE2 | $6.1 \times 10^{+5} \pm 1.0 \times 10^{+4}$ | $4.0 \times 10^{-5} \pm 2.3 \times 10^{-7}$ |
| | CR3022 | $1.0 \times 10^{+6} \pm 5.4 \times 10^{+4}$ | $2.3 \times 10^{-6} \pm 1.0 \times 10^{-6}$ |
| | S309 | $5.0 \times 10^{+5} \pm 7.8 \times 10^{+4}$ | $1.0 \times 10^{-5} \pm 2.4 \times 10^{-6}$ |
| 14 | ACE2 | $5.8 \times 10^{+5} \pm 1.8 \times 10^{+3}$ | $1.1 \times 10^{-4} \pm 2.6 \times 10^{-6}$ |
| | CR3022 | $2.1 \times 10^{+5} \pm 4.4 \times 10^{+3}$ | $2.4 \times 10^{-5} \pm 1.1 \times 10^{-5}$ |
| | S309 | $1.5 \times 10^{+5} \pm 3.2 \times 10^{+3}$ | $1.0 \times 10^{-5} \pm 2.0 \times 10^{-6}$ |

**Table S1: Kinetic parameters for the binding of hACE2, and RBD targeting antibodies CR3022 and S309 to selected SARS-CoV-2 S designs.** Shown are the average and standard deviation for triplicate measurements.

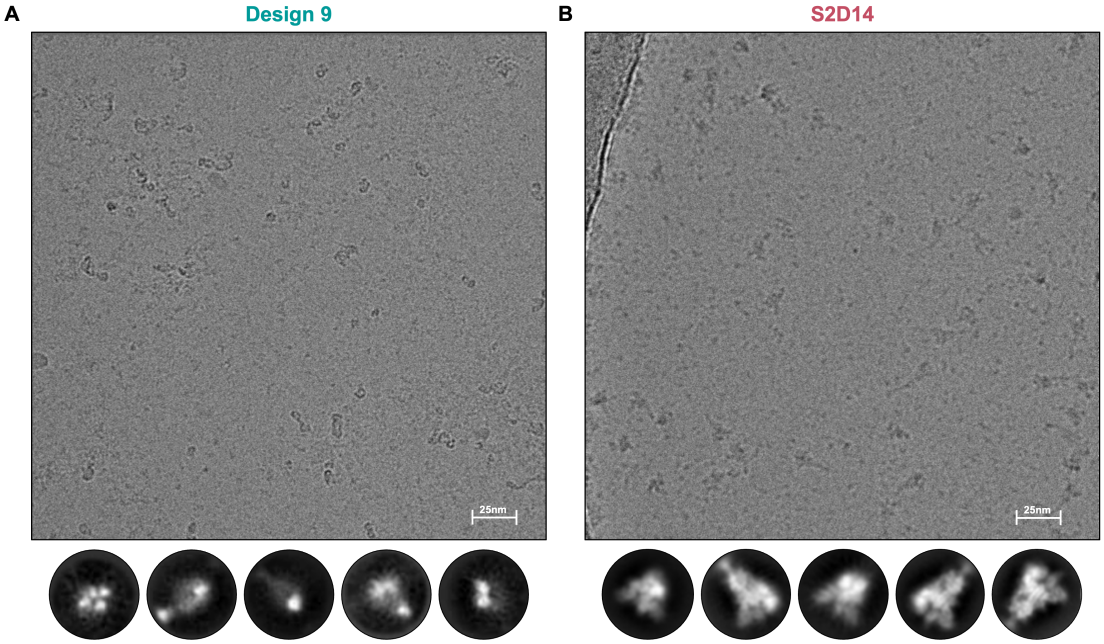


**Fig. S3: Cryo-EM screening of design 9 and S2D14 (design 14) confirms that only S2D14 forms stable prefusion S trimers.** **(A)** Micrograph and 2D class averages (below) of NTD + S2 design 9 illustrating the lack of observable S protein trimers. **(B)** In contrast, images collected for S2D14 contained particles that were consistent with the expected morphology and size of S trimers. 2D classes of S2D14 from Fig.2 are shown below for comparison. Scale bar = 25 nm

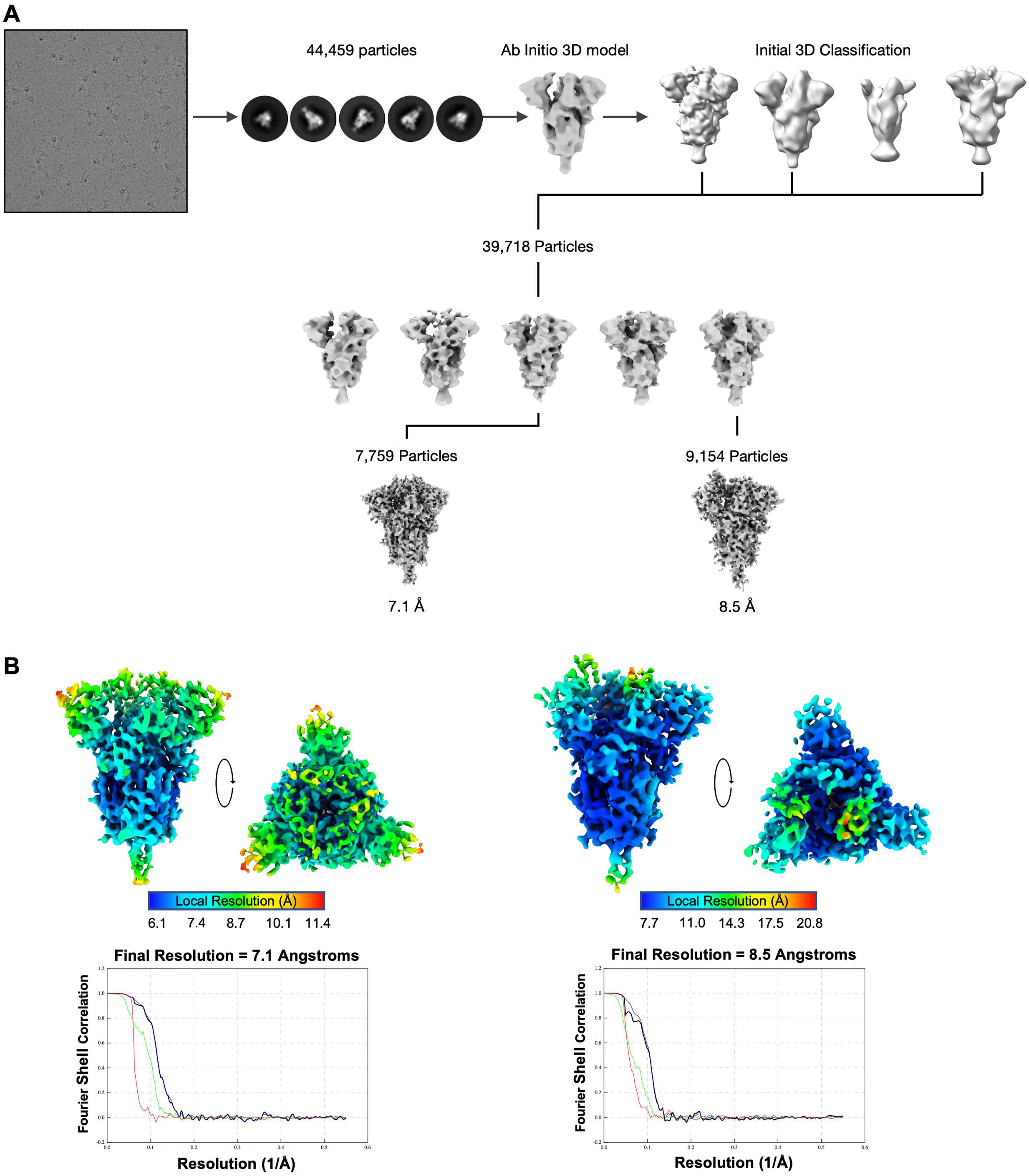


**Fig. S4**: **Single particle reconstruction workflow. (A)** Schematic illustrating the cryo-EM data processing workflow for images collected on a Glacios TEM and using RELION 3.1 image processing suite. **(B)** Local resolution plots were generated for each map using RELION 3.1 and the FSC curves (below) report the resolution using the gold-standard FSC at 0.143 criteria.

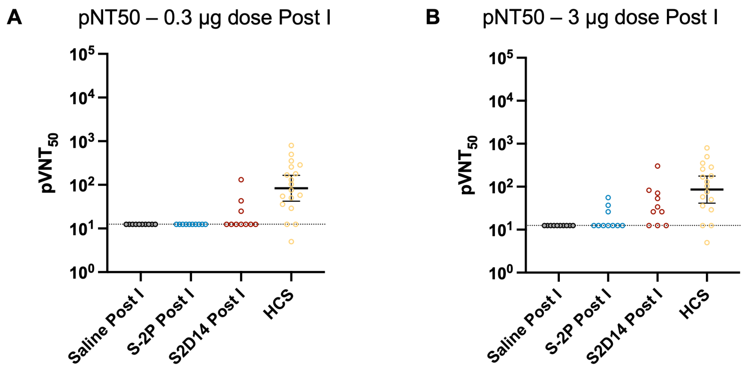


**Fig S5**. **Neutralizing antibody titers from immunized mice against Wuhan post-I compared to human convalescent serum.** Neutralizing antibody titers against the Wuhan strain were analyzed 2 weeks post-I vaccination at two antigen dosages, 0.3 µg **(A)** and 3.0 µg **(B)**. Neither S-2P or S2D14 elicited a neutralizing response post-I compared to human convalescent sera (HCS). For HCS samples, GMT and 95% CI were calculated separately from vaccination groups in Prism GraphPad. See Methods for a description of the sera panel.

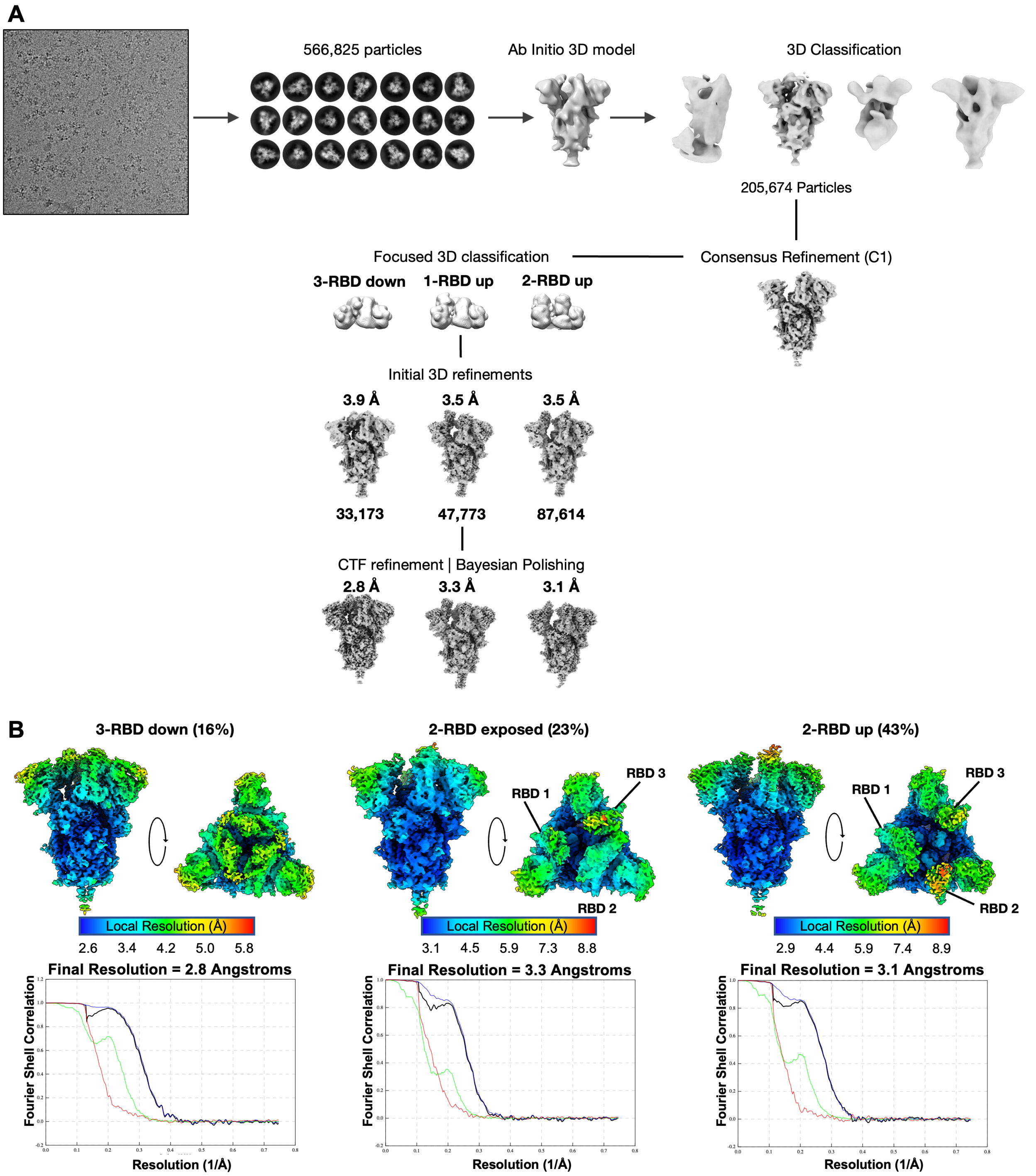


**Fig. S6**: **High-resolution single particle reconstruction workflow. (A)** Schematic illustrating the cryo-EM data processing workflow for images collected on a Titan Krios and using RELION 3.1 image processing suite. **(B)** Local resolution plots were generated for each map using RELION 3.1 and the FSC curves (below) report the resolution using the gold-standard FSC at 0.143 criteria.

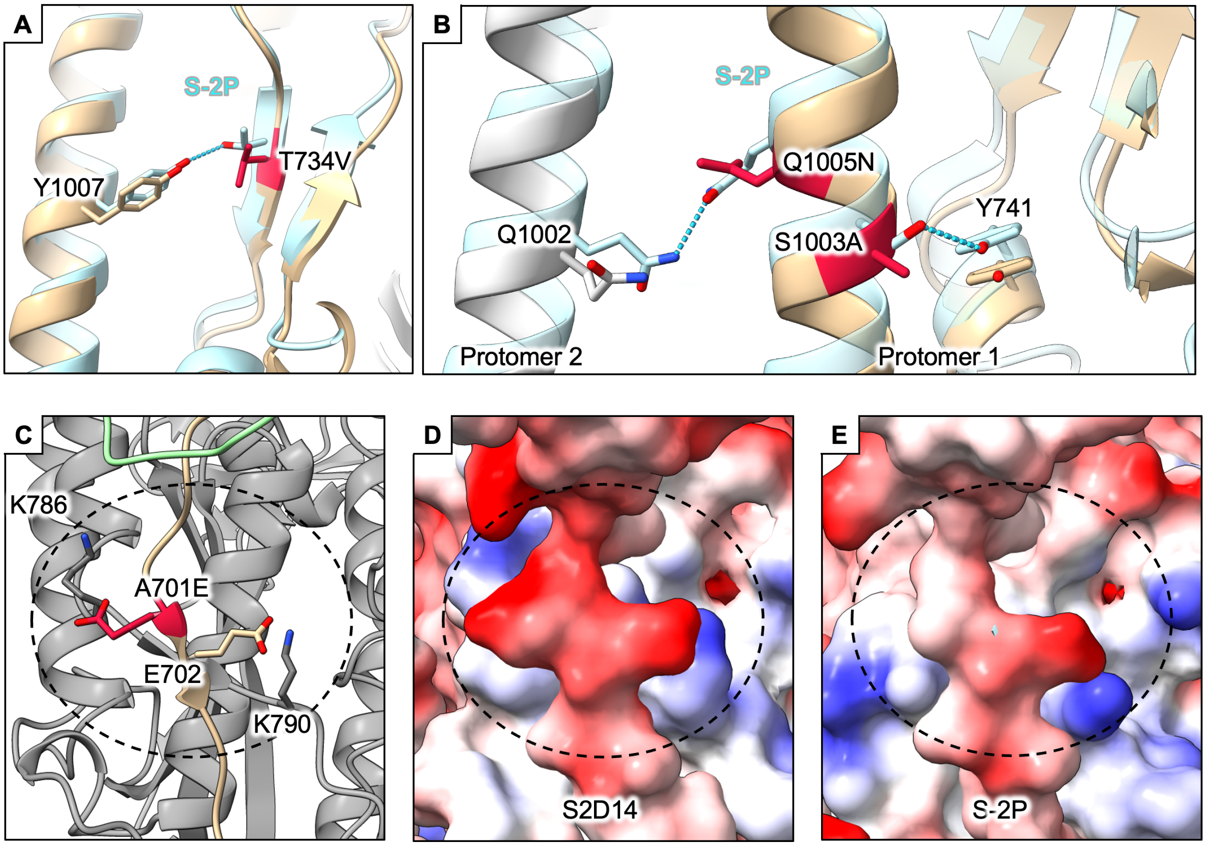


**Fig. S7**: **Mutations within S2D14 affecting hydrogen bond formation and electrostatic interactions.** Hydrogen bonds in S-2P are removed by mutations such as **(A)** T734V, and **(B)** S1003A and Q1005N**.** Hydrogen bonds are depicted as blue dotted lines. **(C)** The A701E mutation in the linker region between the S1 and S2 domain is in proximity to a positively charged lysine at position K786 on a neighboring protomer and adjacent to a salt bridge formed between E702 and K790. Comparison of the S2D14 electrostatic potential **(D)** of the S1/S2 linker to S-2P **(E)** shows a greater negative patch for S2D14 that may strengthen interprotomer contacts via complementary electrostatic interactions.

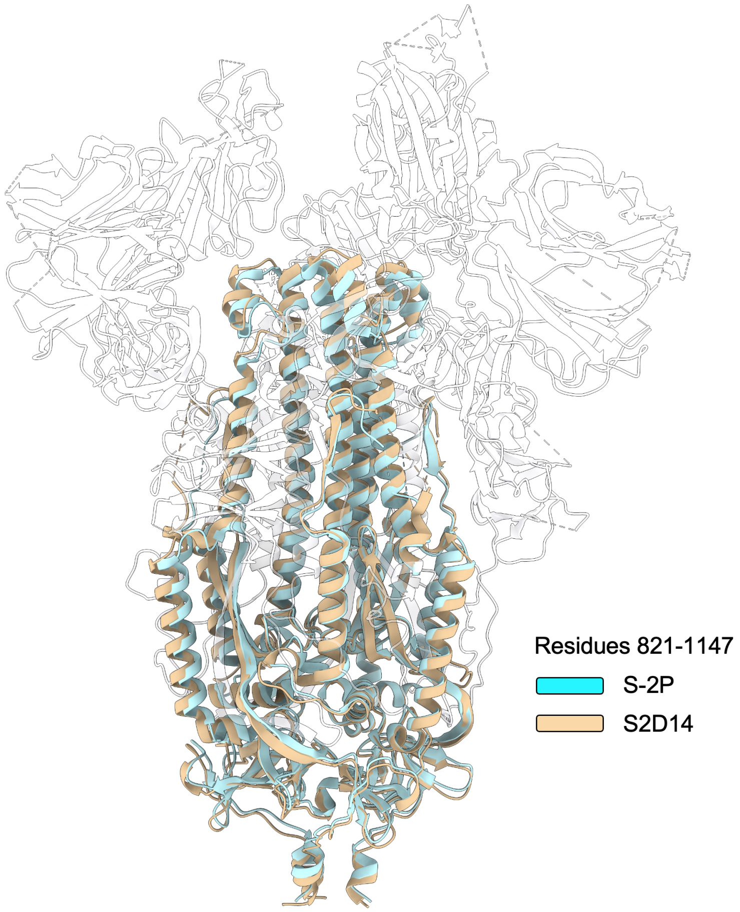


**Fig. S8: S2 domain architecture for S2D14 is not disrupted by S2 domain mutations.** Superposition of S-2P (PDB 6VSB) and S2D14 (two RBD open structure) S2 domain residues 821-1147 reveals an RMSD of 1.6 Å over all $C_{\alpha}$ atoms. S-2P is colored in light blue and S2D14 is colored in tan. The S1 domain of S2D14 is shown as a silhouette for clarity.

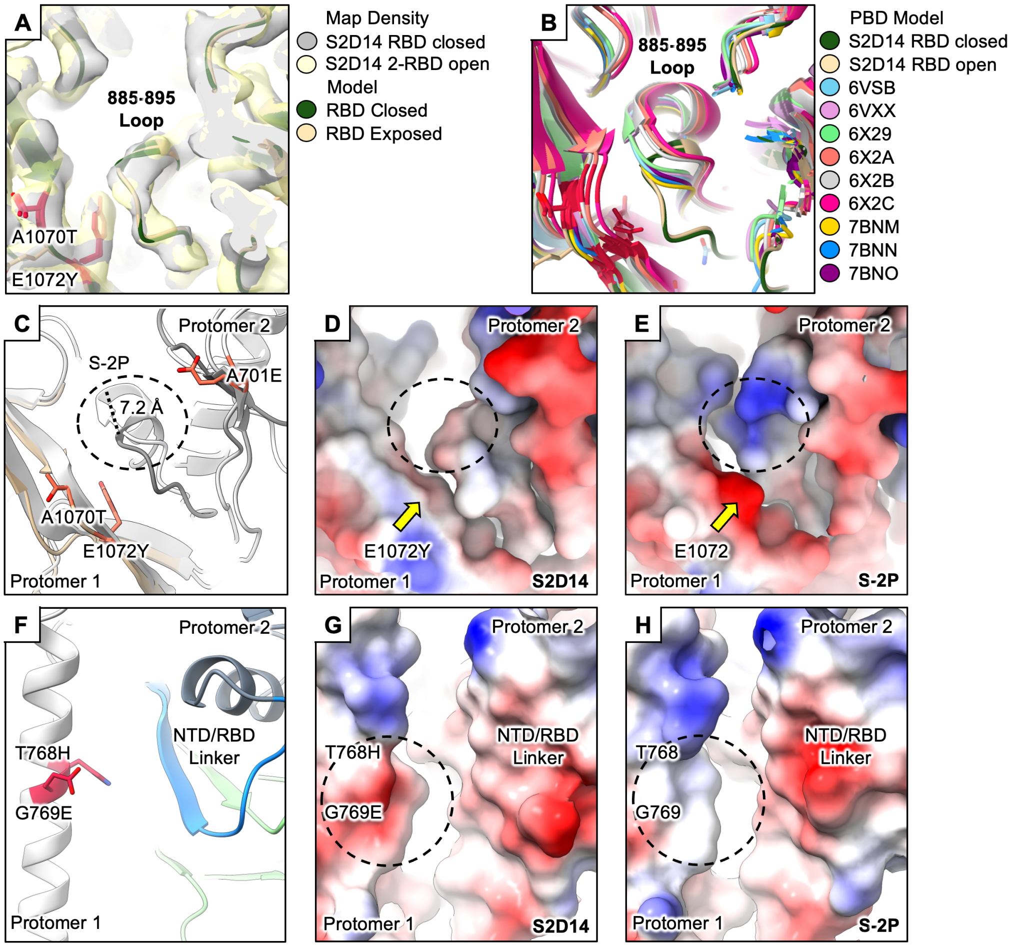


**Fig. S9**: **Mutations within S2D14 that may promote sampling of RBD exposed states. (A)** Superposition of S2D14 structures and EM densities for the three RBD closed and two RBD open state show identical positioning of the 885-895 loop in proximity to the A1070T and T1072Y mutations. **(B)** The position of the 885-895 loop compared with several published structures shows that this loop placement is unique to S2D14. **(C)** Superposition of S-2P (light grey) and S2D14 shows an ~7.2 Å shift of the 885-895 loop towards the E1072Y mutation. **(D)** The same orientation as shown in (C) but with S2D14 depicted in electrostatic surface. The dotted circle highlights a gap in the S2D14 trimer interior caused by displacement of the 885-895 loop, which is now packed against a hydrophobic patch created by the E1072Y mutation (indicated by a yellow arrow). **(E)** The same orientation as shown in (C) but with S-2P depicted in electrostatic surface. The dotted circle highlights the position of the 885-895 loop towards the S-2P trimer interior. The wild-type E1072 is indicated by a yellow arrow. **(F)** Structure of the T768H and G769E mutations in S2D14 in proximity to the NTD/RBD linker region on an adjacent protomer which acts as part of the hinge for RBD opening. **(G)** The same orientation as shown in (F) but with S2D14 depicted in electrostatic surface. The T768H and G769E mutation imparts a negative surface potential (indicated by a dashed circle) that may destabilize the NTD/RBD linker region and promote RBD opening. **(H)** The same orientation as shown in (F) but with S-2P depicted in electrostatic surface. The wild-type T768 and G769 residues creates a more neutral surface potential compared to S2D14 (indicated by a dashed circle).

1. Troshin, P.V., J.B. Procter, and G.J. Barton, *Java bioinformatics analysis web services for multiple sequence alignment--JABAWS:MSA.* Bioinformatics, 2011. **27**(14): p. 2001-2.
